# A viral toolbox for conditional and transneuronal gene expression in zebrafish

**DOI:** 10.1101/2021.03.25.436574

**Authors:** Chie Satou, Rachael L. Neve, Hassana K. Oyibo, Pawel Zmarz, Kuo-Hua Huang, Estelle Arn Bouldoires, Takuma Mori, Shin-ichi Higashijima, Georg B. Keller, Rainer W. Friedrich

**Author notes:** **Contact:** Rainer Friedrich, Friedrich Miescher Institute for Biomedical Research, Maulbeerstrasse 66, 4058 Basel, Switzerland.

## Abstract

The zebrafish is an important model in systems neuroscience but a key limitation is the lack of viral tools to dissect the structure and function of neuronal circuitry. We developed methods for efficient gene transfer and retrograde tracing in adult and larval zebrafish by herpes simplex viruses (HSV1). HSV1 can be combined with the Gal4/UAS system to target cell types with high spatial, temporal and molecular specificity. We also established methods for efficient transneuronal tracing by modified rabies viruses in zebrafish. We demonstrate that HSV1 and rabies viruses can be used to visualize and manipulate genetically or anatomically identified neurons within and across different brain areas of adult and larval zebrafish. An expandable library of viruses is provided to express fluorescent proteins, calcium indicators, optogenetic probes, toxins and other molecular tools. This toolbox creates new opportunities to interrogate neuronal circuits in zebrafish through combinations of genetic and viral approaches.

## Introduction

The zebrafish is an important vertebrate model in systems neuroscience because its small, optically accessible brain provides unique opportunities to analyze neuronal circuits and behavior^1,2^. Key methods established in zebrafish include large-scale imaging of neuronal population activity, behavioral approaches including virtual realities, and genetic manipulations^1–8^, but efficient methods for viral gene transfer are still lacking. Viral vectors are important experimental tools in mammals because they enable the visualization and manipulation of defined neurons with high spatial, temporal and molecular specificity, and because they can bypass the need to generate transgenic animals^9,10^. Moreover, engineered viral tools that cross synapses allow for the visualization and physiological analysis of synaptically connected neuronal cohorts^11–13^. Our goal was to establish similar methods for specific viral gene transfer and transneuronal tracing in zebrafish.

## Results

### Optimization of the temperature regime for viral gene transfer

Previous studies demonstrated that adeno-associated viruses (AAVs), which are widely used for gene transfer in mammals and other amniotes, fail to infect neurons in the zebrafish brain^14^. Infection of zebrafish neurons by other viruses has been reported but efficiency was usually low and viral vectors for conditional gene expression in transgenic fish have not been described. To improve upon these points, we first focused on herpes simplex virus 1 (HSV1), a DNA virus that can infect zebrafish neurons both locally and retrogradely via projecting axons without obvious signs of cytotoxicity^15^. We first explored whether HSV1-mediated gene transfer can be further improved by optimizing the temperature regime. Zebrafish are usually kept at 26 – 28.5 °C but the temperature range of natural habitats is broad (up to >38 °C) and temperature tolerance in the laboratory extends up to ∼41 °C^16,17^. We therefore tested whether viral gene expression is more efficient at temperatures near those of mammalian hosts (37 °C).

After injection of amplicon type HSV1 viruses into the brain of adult or larval zebrafish using established procedures^15^, swimming behavior appeared normal both at standard laboratory temperatures (26 - 28.5 °C) and when fish were kept at 35 – 37 °C (Supplementary Movies 1 – 3). To further examine effects of temperature on behavior we trained two groups of adult zebrafish in an odor discrimination task^6,18^ that comprised one day of acclimatization to the setup followed by five days of appetitive conditioning (nine training trials with a rewarded odor [CS+] and with a non-rewarded odor [CS-] each per day). Group 1 (control) underwent no surgery and was kept at the standard laboratory temperature. Group 2 was injected with an HSV1 and subsequently kept at 36 °C for two days before training commenced (Fig. S1a). Learning was assessed by a standard discrimination score and not significantly different between groups. These results confirm that swimming behavior and olfactory discrimination learning were not significantly impaired by virus injection or subsequent incubation at temperatures near 37 °C.

To examine the temperature-dependence of HSV1-mediated gene expression we injected adult zebrafish expressing green fluorescent protein (GFP) under the *vglut1* promoter (Tg[*vglut1*:GFP]) with HSV1[*LTCMV*:DsRed], an HSV1 with an insert encoding the red-fluorescent protein DsRed under the control of a non-specific promoter for long-term expression (*LTCMV*). Injections were targeted unilaterally into the olfactory bulb (OB) and fish were then maintained at 26 °C or 36 °C for 6 days after injection (Fig. 1a). Consistent with previous results^15^, reporter expression (DsRed) was observed in the injected OB (Fig. S1c). Moreover, retrogradely labeled neurons were present in telencephalic areas that project to the OB, most notably in the posterior zone of the dorsal telencephalon (Dp), the homolog of mammalian olfactory cortex (Fig. S1c). After incubation at 36 °C, substantially more neurons expressed DsRed (Fig. 1b). In addition, retrograde labeling was detected in a small but distinct population of neurons in the dorsal telencephalon that was not seen at 26 °C (Fig. 1b, c). DsRed expression at 36 °C was first detected ∼12 hours after injection and stable for at least 10 days (Fig. S1d). Moreover, increasing the temperature from 26 °C to 36 °C three days after the injection failed to enhance expression (Fig. 1a, c). These results show that adjusting the temperature to that of mammalian hosts can substantially increase the efficiency of HSV1-mediated viral gene transfer in zebrafish.

**Fig 1.**
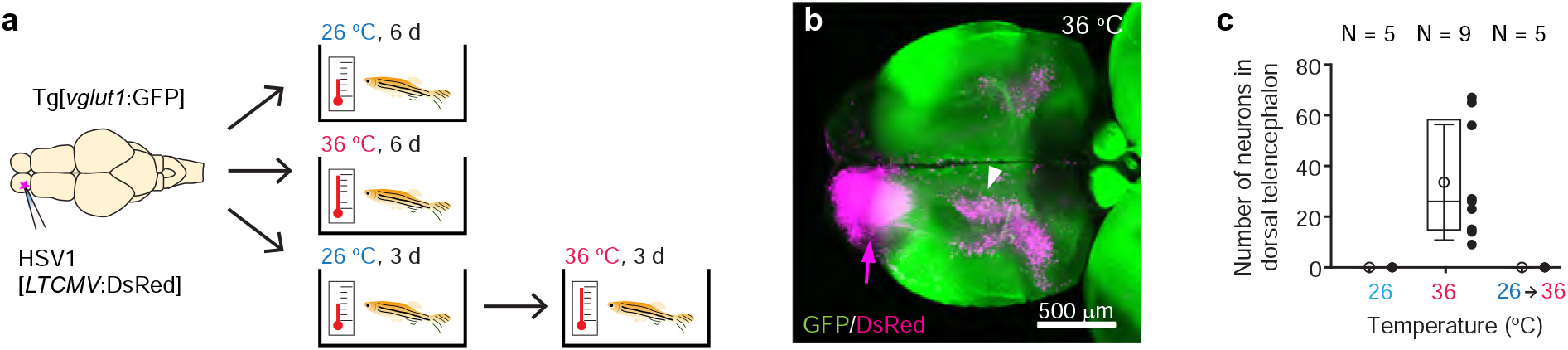
HSV1-mediated gene delivery in adult zebrafish. **(a)** Procedure to test temperature-dependence of HSV1-mediated gene expression. **(b)** Maximum intensity projection after injection of HSV1[*LTCMV*:DsRed] into one OB (arrow) of a Tg[*vglut1*:GFP] fish and incubation at 36 °C. White arrowhead indicates the OB-projecting area in the dorsal telencephalon used for quantification in Fig. 1c and Fig. 2f**(f)**. *vglut1*:GFP expression served as a morphological marker. **(c)** Mean number of labeled neurons in the dorsal telencephalon after injection of HSV1[*LTCMV*:DsRed] into the ipsilateral olfactory bulb and incubation at different temperatures. In this and similar plots, black dots represent data from individual fish, box plot indicates median and 25^th^ and 75^th^ percentiles, circles and error bars indicate mean and s.d., respectively, over individual fish. N: number of fish.

To explore viral gene delivery at earlier developmental stages we injected HSV1[*LTCMV*:GFP] into the optic tectum of zebrafish larvae at 3 days post fertilization (dpf) and examined expression 48 h later. Consistent with previous observations^15^, expression of GFP was observed when fish were kept at 28.5 °C (N = 3 fish) but the number of GFP-positive cells further increased when temperature was raised to 32 °C (N = 5) or 35 °C (N = 10; Fig. S2a). Strong and widespread expression was also observed when the virus was injected at 5 dpf (N = 14) or 14 dpf (N = 9) and fish were subsequently kept at 35 °C for 48 h (Fig. S2b,c). When HSV1[*LTCMV*:GFP] was injected into muscles of the trunk at 5 dpf, strong and selective retrograde labeling was observed in motor neurons in the spinal cord after keeping fish at 35 °C for 48 h (N = 4; Fig. S2d). These results indicate that HSV1 can be used for gene delivery and retrograde neuronal tracing throughout development.

### Intersection of HSV1 with the Gal4/UAS system

Viral vectors are often combined with transgenic lines to target specific cell types using two-component expression systems (e.g., injection of a Cre-dependent viral construct into Cre-expressing mice). In zebrafish, the most widely used two-component expression system is the Gal4/UAS system. We therefore explored the possibility to combine viral delivery of UAS-dependent expression constructs with Gal4-expressing driver lines. We first created a Gateway expression vector^20^ to simplify the construction of HSV1 for UAS-dependent expression (Fig. 2a). We then generated HSV1[*UAS*:TVA-mCherry] to expresses a fusion of the transmembrane protein TVA and the red fluorescent protein mCherry under UAS control. This virus did not drive expression when injected into the brain of wt zebrafish (N = 3; not shown).

**Fig 2.**
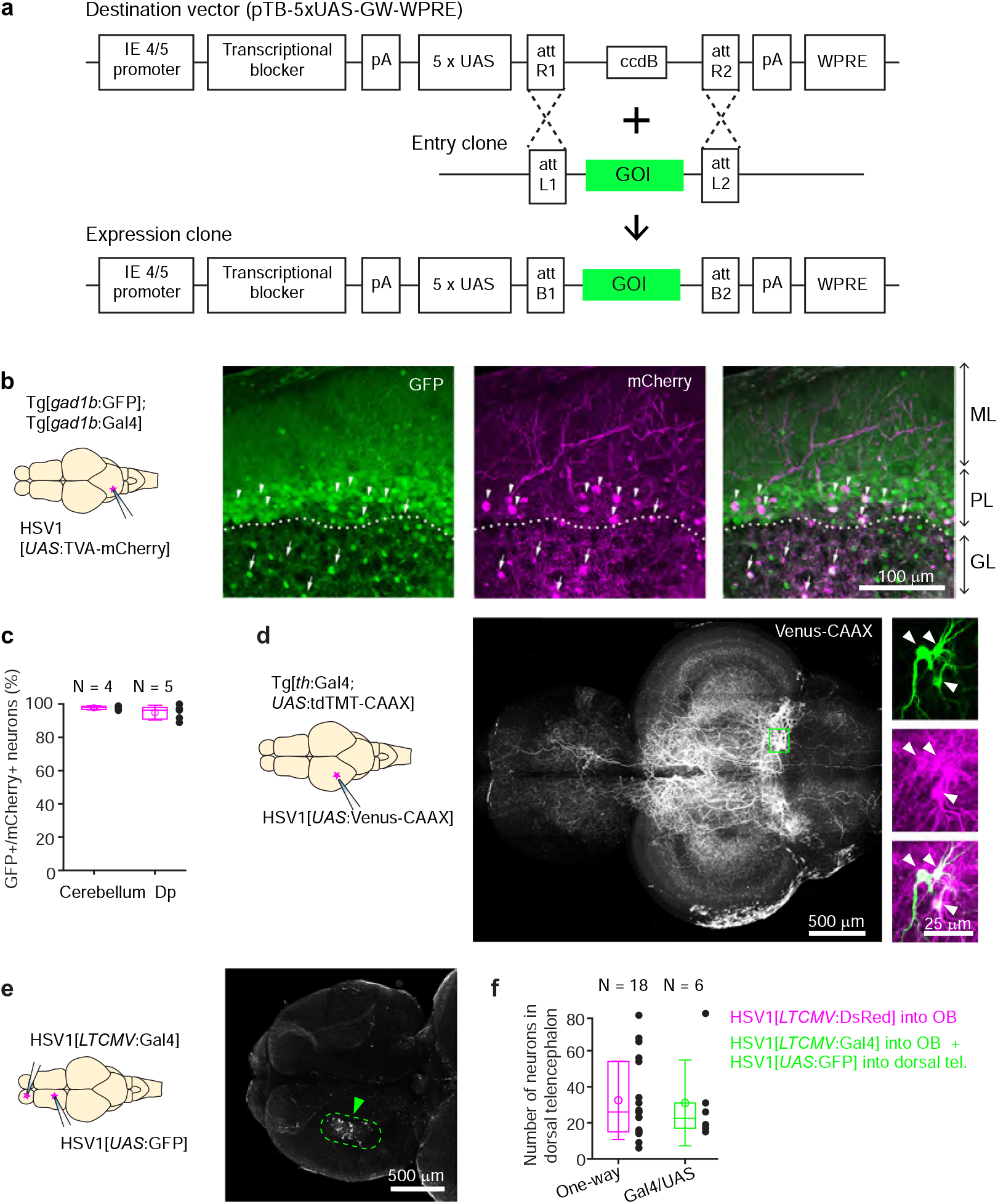
Conditional HSV1-mediated gene expression using the Gal4/UAS system. **(a)** Construction of the UAS vector for HSV packaging. Genes of interest (GOI) are inserted downstream of the 5xUAS sequences by recombination cloning using the Gateway system. The transcriptional blocker minimizes leaky expression^60^. **(b)** Injection of HSV1[*UAS*:TVA-mCherry] into the cerebellum of Tg[*gad1b*:Gal4; *gad1b*:GFP] double transgenic fish. Note co-localization of mCherry and GFP in Purkinje cells (arrowheads) and putative Golgi cells (arrows). ML: molecular layer; PL: Purkinje layer; GL: granular layer. **(c)** Fraction of mCherry-positive neurons that co-expressed GFP after injection of HSV1[*UAS*:TVA-mCherry] into the cerebellum or Dp of Tg[*gad1b*:Gal4; *gad1b*:GFP] double transgenic fish. N: number of fish. **(d)** Injection of HSV1[*UAS*:Venus-CAAX] into the optic tectum of Tg[*th*:Gal4; *UAS*:tdTomato-CAAX] fish. Venus-CAAX was expressed by a small number of neurons with somata in the locus coeruleus and extensive projections to the optic tectum and other brain areas. Images on the right are close-ups of the boxed region showing co-expression of Venus-CAAX (green) with tdTomato (red) in the locus coeruleus. **(e)** Injection of HSV1[*LTCMV*:Gal4] into the OB and HSV1[*UAS*:GFP] into the dorsal telencephalon of wt fish. Note selective expression of GFP in OB-projecting neurons (arrowhead; dashed outline). **(f)** Number of neurons labeled in the dorsal telencephalon by a single injection of HSV1[*LTCMV*:DsRed] into the OB (‘One way’) or by two injections using the two-component Gal4/UAS system (‘Gal4/UAS’). N: number of fish.

We injected HSV1[*UAS*:TVA-mCherry] into the cerebellum of adult Tg[*gad1b*:GFP; *gad1b*:Gal4] fish^18^, which express Gal4 and GFP under the control of the *gad1b* promoter, a marker of GABAergic neurons. Consistent with the distribution of GABAergic neurons in the cerebellum, GFP was expressed in Purkinje neurons and in a sparse neuronal population in the granular layer, presumably Golgi cells. mCherry was co-expressed with GFP in a subset of Purkinje neurons and putative Golgi cells (97.7 ± 1.2 % of mCherry-positive neurons were also GFP-positive; mean ± s.d.; N = 4 fish; Fig. 2b,c). No expression was detected in the dense population of GFP-negative granule cells in the granular layer.

Co-expression of mCherry and GFP was observed also when the injection of HSV1[*UAS*:TVA-mCherry] was directed at a cluster of GABAergic neurons near Dp in adult Tg[*gad1b*:GFP; *gad1b*:Gal4] fish (94.8 ± 4.5 % of mCherry+ neurons coexpressed GFP; mean ± s.d.; N = 5 fish; Fig. 2c, Fig. S3). Interestingly, mCherry-expressing neurites arborized extensively in the lateral pallium but spared a region of Dp immediately adjacent to the *gad1b*-positive cluster, indicating that the dorsal telencephalon contains specific long-range projections of GABAergic neurons (Fig. S3a). In zebrafish larvae (7 dpf) we injected HSV1[*UAS*:GFP] into the hindbrain of Tg[*gad1b*:Gal4; *gad1b*:DsRed] fish (N = 3) and detected expression of GFP selectively in DsRed-positive cells in the hindbrain and cerebellum (Fig. S2e). These observations indicate that HSV1 can be combined with the Gal4/UAS system to enhance the specificity of cellular targeting.

To corroborate this conclusion we designed experiments to target sparse neuronal populations. We first injected HSV1[*UAS*:Venus-CAAX] into the optic tectum of Tg[*th*:Gal4; *UAS*:tdTomato-CAAX] fish, which express Gal4 and the red-fluorescent protein tdTomato-CAAX under the control of the tyrosine hydroxylase-1 promoter, a marker for catecholaminergic neurons. This procedure resulted in the selective expression of the membrane-associated protein Venus-CAAX in a small number of tdTomato-CAAX-positive neurons in the locus coeruleus with complex long-distance axonal projections, indicating that expression was specifically directed to catecholaminergic (noradrenergic) neurons (Fig. 2d; N = 4 fish). In additional experiments we first injected HSV1[*LTCMV*:Gal4] into the OB of wt zebrafish and subsequently injected HSV1[*UAS*:GFP] into the dorsal telencephalon. Expression of GFP was observed specifically in OB-projecting neurons of the dorsal telencephalon without expression elsewhere (Fig. 2e). To assess the efficiency of this intersectional targeting approach we compared the number of labeled neurons using either the Gal4/UAS system or the one-component approach (injection of HSV1[*LTCMV*:DsRed] into OB). Both approaches yielded similar numbers of labeled neurons in the dorsal telencephalon (Fig. 2f). We therefore conclude that HSV1 can be combined with the Gal4/UAS system in intersectional approaches with high efficiency and low leakiness.

We next explored strategies for co-expression of multiple transgenes in the same neurons. When red and green reporter constructs were co-packaged into the same virus (HSV1[*UAS*:TVA-mCherry & *UAS*:GFP]), injection into the cerebellum of Tg[*gad1b*:Gal4] resulted in a high rate of co-expression, as expected (89.5 ± 8.7 % co-expression; mean ± s.d.; N = 3 fish; Fig. S4a, c). However, the rate of co-expression did not reach 100 %, possibly because not all virus particles received both constructs during packaging. Interestingly, co-injection of the same responder constructs packaged into separate viruses (HSV1[*UAS*:TVA-mCherry] and HSV1[*UAS*:GFP]) produced a similar rate of co-expression, even when overall expression was sparse (84.3 ± 6.0 % co-expression; mean ± s.d.; N = 3 fish; Fig. S4b, c). These results indicate that co-packaging and co-injection of viruses can be used to express multiple transgenes in the same neurons.

We also tested HSV1 as a tool for functional manipulations of neurons and behavior. We first injected HSV1[*LTCMV*:Gal4] into the OB of wt zebrafish and subsequently co-injected HSV1[*UAS*:GCaMP6f] and HSV1[*UAS*:Chrimson-tdTomato] into the dorsal telencephalon. This procedure resulted in the co-expression of the green-fluorescent calcium indicator GCaMP6f ^21^ and the red-light-gated cation channel Chrimson-tdTomato^22^ in OB-projecting neurons of the dorsal telencephalon. Brief (500 ms) illumination with red light evoked GCaMP6f fluorescence transients that increased in amplitude with increasing light intensity (Fig. S5; 11 neurons in N = 1 fish). Hence, transgene-expressing neurons were functional and responsive to optogenetic stimulation. To explore strategies for neuronal ablations we expressed tetanus toxin light chain (TeNT) in GABAergic neurons of the cerebellum by injecting HSV1[*UAS*:TeNT-GFP] into the cerebellum of Tg[*gad1b*:Gal4] fish. Injected fish showed abnormal body posture and swimming patterns while control fish injected with HSV1[*UAS*:GFP] swam normally (Supplementary Movie 4). HSV1-mediated gene transfer thus offers a wide range of opportunities for anatomical and functional experiments in zebrafish.

We have so far assembled a collection of 26 different HSV1s for applications in zebrafish (Supplementary Table1). 15 of these drive expression of fluorescent markers, calcium indicators, Gal4 or optogenetic probes under the control of the non-specific LTCMV promoter for regional transgene expression and retrograde tracing. The remaining 11 HSV1s drive expression of fluorescent markers, optogenetic probes, toxins and other proteins under UAS control for intersectional targeting of neurons using the Gal4/UAS system. This toolbox is available and can be further expanded using our UAS-containing expression vector (Fig. 2a).

### Transneuronal viral tracing in zebrafish

The ability of some viruses to cross synapses has been exploited to express transgenes in synaptically connected cohorts of neurons. In zebrafish and other species, neurons were traced across one or multiple synapses using engineered vesicular stomatitis viruses (VSVs) but these tools have not been used widely as biological tools, possibly because they can exhibit toxicity^23–25^. In rodents, modified rabies viruses have become important tools to analyze connectivity and structure-function relationships in neuronal circuits. These vectors can infect specific “starter” neurons and are transmitted retrogradely to presynaptic neurons across one synapse. Specific infection is achieved by expressing the receptor protein TVA in the starter neuron and pseudotyping the virus with the envelope protein EnvA^11^. To limit retrograde transfer to one synapse, an essential glycoprotein (G) is deleted from the viral genome and supplied in *trans* only in the starter neurons. In zebrafish, the G-deleted rabies virus (RVΔG) has been reported to infect neurons^14^ and to cross synapses when complemented with G but the efficiency of retrograde synaptic transfer appeared very low^26^.

To enhance the efficiency of viral infection and transneuronal tracing we first explored the effect of temperature. When we injected EnvA-coated RVΔG expressing GFP (EnvA-RVΔG-GFP) into the telencephalon of wt fish at 36 °C, no GFP expression was observed (Fig. S6). We then injected EnvA-RVΔG-GFP into the telencephalon of Tg[*gad1b*:Gal4; *UAS*:TVA-mCherry] fish that express TVA-mCherry in GABAergic neurons. After 6 days, efficient and selective infection of neurons expressing GFP was observed at 36 °C but not at 26 °C, nor when the temperature was raised from 26 °C to 36 °C three days post injection (Fig. S7). Hence, infection of zebrafish neurons by EnvA-RVΔG-GFP required TVA and was substantially more efficient when the temperature was close to the body temperature of natural hosts. Subsequent experiments were therefore performed at 35 – 37 °C.

To assess potential toxicity of RVΔG we injected EnvA-RVΔG-GFP into the OB of Tg[*gad1b*:Gal4; *UAS*:TVA-mCherry] fish as before, dissociated the OB, and performed RNA sequencing after fluorescence-activated cell sorting. This approach allowed us to compare transcriptomes of GABAergic neurons infected by RVΔG (green and red) to transcriptomes of non-infected GABAergic neurons expressing only the TVA receptor (red only; Fig. S8). Of 19’819 endogenous genes analyzed, 522 were significantly downregulated while 27 were significantly upregulated in infected cells. Among these, stress markers occurred with approximately average frequency (downregulated: 27 out of 471 stress marker genes; upregulated: 1/471) and cell death markers were underrepresented (downregulated: 14/651 genes; upregulated: 4/651; Fig. 3a-c; Supplementary Table2). The 65 gene ontology (GO) terms that were significantly associated with the set of regulated genes were primarily related to immune responses while GO terms related to stress, cell death, electrophysiological properties or synapses were rare or absent (Supplementary Table 3). These results indicate that RVΔG does not have major effects on the health or physiological status of infected neurons.

**Fig 3.**
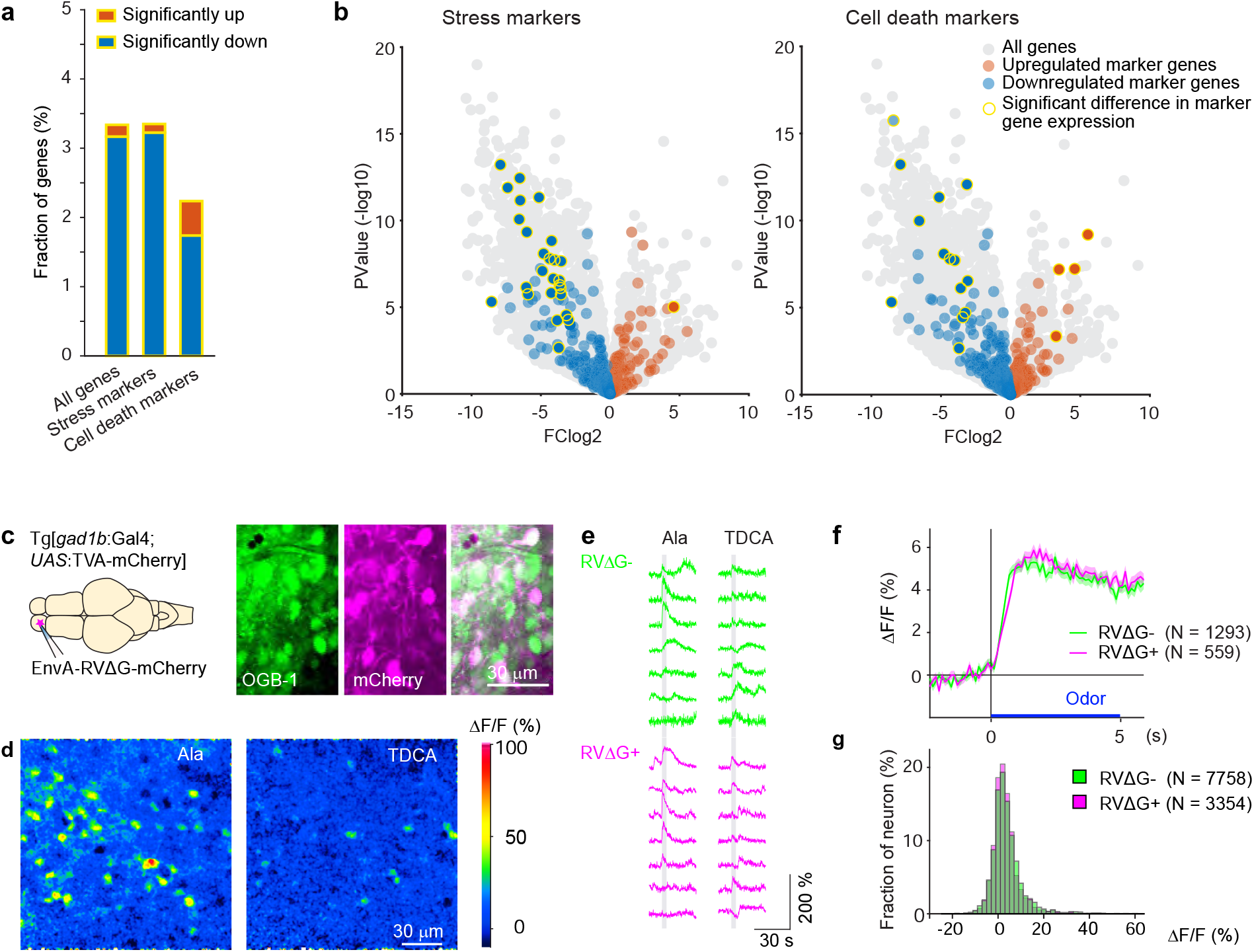
Effects of RV infection on neuronal health and function. **(a)** Fraction of genes that were significantly up- or down-regulated genes in RVΔG-infected cells out of all 19’819 genes, out of the 471 stress markers (GO:0033554), and out of the 651 cell-death markers (GO:0008219). Differences in expression level were considered significant when abs(logFC (fold change)) >3, log(counts per million reads mapped) >3, and FDR<0.05. The FDR (False Discovery Rate) corrects for multiple testing. **(b)** Volcano plots displaying differential gene expression in RVΔG-infected and uninfected cells. Colored dots indicate stress markers (left) and cell death markers (right) (orange: upregulated, blue: downregulated). Yellow outline depicts statistically significant difference in expression level. **(c)** OGB-1 labeling and mCherry expression in the deep (granule cell) layer of the adult zebrafish OB after injection of EnvA-RVΔG-mCherry into the OB of Tg[*gad1b*:Gal4; *UAS*:TVA-mCherry] fish and bolus loading of OGB-1. **(d)** Ca^2+^ signals evoked by two different odors in the same optical plane (single trials). Odors: alanine (Ala), taurodeoxycholic acid (TDCA). **(e)** Randomly selected responses of seven infected (magenta) neurons and seven uninfected (green) neurons from the same optical plane to two odors (single trials). **(f)** Odor-evoked Ca^2+^ signals of infected (N = 559) and uninfected (N = 1293) OB cells from *N* = 4 fish, averaged over all odors (N = 6) and repetitions (N = 3 for each odor). Shading indicates s.e.m.; bar indicates odor stimulation. **(g)** Distribution of response amplitudes in non-infected and infected cells to different odors (N = 6), averaged over trials (N = 3). Distributions of were not significantly different (p = 0.24, Kolmogorov–Smirnov test).

To directly compare neuronal activity of infected and uninfected neurons we targeted the dense population of GABAergic interneurons in the deep layers of the OB. We injected EnvA-RVΔG-mCherry into the OB of adult Tg[*gad1b*:Gal4; *UAS*:TVA-mCherry] fish and detected infection in a subset of neurons by the strong cytoplasmic and nuclear expression of mCherry, which could easily be distinguished from the weak, membrane-associated background expression of TVA-mCherry (Fig. 3c). We then loaded neurons non-specifically with the green-fluorescent calcium indicator Oregon Green 488 BAPTA-1 (OGB1) by bolus injection of the AM ester^27^ and measured odor responses of all neurons simultaneously (Fig. 3d). No obvious differences were detected in the time course or amplitude distribution of odor responses between infected neurons (N = 1293 neurons from four fish) and uninfected neurons (N = 559 from the same four fish; Fig. 3e-g). Together, these results indicate that infection with RVΔG did not compromise the health or physiological function of neurons.

Efficient transneuronal spread of the rabies virus depends on the expression level of the viral glycoprotein in starter cells^28,29^. We therefore took two steps to enhance glycoprotein expression. First, we optimized codon usage for zebrafish. Second, we expressed TVA and the codon-optimized glycoprotein (zoSADG) using HSV1 because viral vectors typically reach higher expression levels than transgenics^15^. In rodents, starter neurons expressing high levels of G often disappear in parallel with the emergence of transneuronal expression, presumably because long-term expression of G is toxic^30,31^. We therefore determined the time course of transgene expression under different experimental conditions. We first focused on the cerebellum where GABAergic Purkinje neurons receive local synaptic input from different types of cerebellar neurons and long-range input from neurons in the contralateral inferior olive (climbing fibers; Fig. 4a). We injected a mixture of HSV1[*UAS*:TVA-mCherry] and EnvA-RVΔG-GFP into the cerebellum of Tg[*gad1b*:Gal4] fish and examined expression for up to 10 days. As observed before (Fig. 2b), mCherry expression was localized to Purkinje neurons and putative Golgi cells. Neurons expressing GFP were concentrated around the injection site and usually co-expressed mCherry (Fig 4b, d). Rarely, GFP expression was observed in mCherry-negative neurons (Fig. 4d), which may indicate TVA-independent viral entry. However, as no GFP expression was observed when EnvA-RVΔG-GFP was injected into wt animals (Fig. S6), it appears more likely that GFP+/mCherry-neurons initially expressed TVA but subsequently lost expression due to the slight decline of expression driven by the viral promoter (Fig. S1b). Co-expression of mCherry and GFP was stable for at least 10 days after injection and GFP+/mCherry-neurons remained very rare (Fig. 4d). These results confirm the specificity of EnvA-RVΔG-GFP infection and provide additional evidence that expression of TVA-mCherry or the infection by RVΔG alone are not toxic in the absence of glycoprotein. No expression of GFP was observed in the inferior olive, consistent with the expectation that RVΔG does not spread in the absence of glycoprotein.

**Fig 4.**
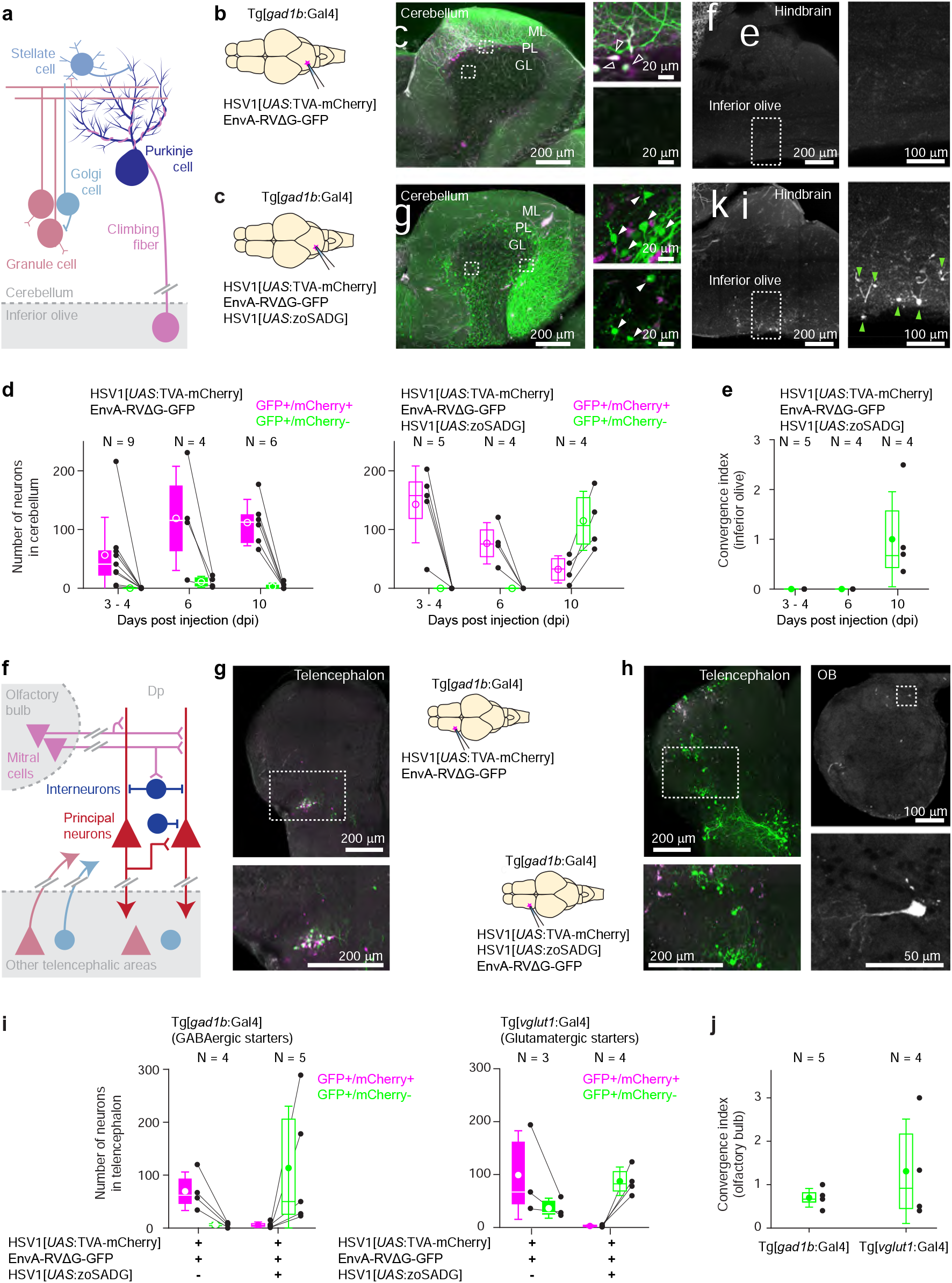
Transneuronal tracing using pseudotyped rabies virus in adult zebrafish. **(a)** Schematic of the cerebellar circuit. Glutamatergic neurons are shown in red colors, GABAergic neurons in blue colors. Purkinje cells receive extra-cerebellar input exclusively from the inferior olive. **(b)** Co-injection of EnvA-RVΔG-GFP and HSV1[*UAS*:TVA-mCherry] into the cerebellum of Tg[*gad1b*:Gal4] fish in the absence of glycoprotein. Left: schematic. Center: expression of TVA-mCherry (magenta) and GFP (green) in the cerebellum. Regions in the Purkinje and granular layers (dashed rectangles) are enlarged. Unfilled white arrowheads indicate GFP+/mCherry+ neurons. Right: expression of GFP in the hindbrain. Region covering the inferior olive (dashed rectangle) is enlarged. Expression of GFP was restricted to putative starter neurons; no expression was detected in the inferior olive. ML: molecular layer; PL: Purkinje layer; GL: granular layer. **(c)** Same as in (b) but the glycoprotein (zoSADG) was supplied to starter neurons in *trans* by co-injection of HSV1[*UAS*:zoSADG]. Filled white arrowheads indicate GFP+/mCherry-neurons. Note expression of GFP in putative granule cells and in neurons of the inferior olive, indicating transneuronal spread. **(d)** Number of neurons that expressed GFP and mCherry (putative starter neurons) or GFP alone (putative presynaptic neurons) at different time points after injection of EnvA-RVΔG and HSV1[*UAS*:TVA-mCherry] into the cerebellum of Tg[*gad1b*:Gal4] fish. Left: without glycoprotein; right: with trans-complementation of zoSADG in starter neurons. Note that labeling of putative presynaptic neurons emerged between 6 and 10 days post injection only when zoSADG was trans-complemented in starter neurons. In all plots, black dots represent data from individual fish, box plot indicates median and the 25^th^ and 75^th^ percentiles, circles and error bars indicate mean and s.d. over individual fish. N: number of fish. **(e)** Convergence index for the projection from the inferior olive to the cerebellum at different time points. The convergence index is the numerical ratio of transneuronally labeled neurons (GFP+/mCherry-neurons in the inferior olive) and putative starter cells in the cerebellum (GFP+/mCherry+ Purkinje neurons). N: number of fish. **(f)** Schematic of the putative circuitry in telencephalic area Dp. Glutamatergic neurons are shown in red colors, GABAergic neurons in blue colors. Long-range projections from mitral cells in the olfactory bulb terminate on glutamatergic neurons and on GABAergic interneurons in Dp. Additional long-range projections originate in other telencephalic areas. **(g)** Co-injection of EnvA-RVΔG and HSV1[*UAS*:TVA-mCherry] into Dp of Tg[*gad1b*:Gal4] fish in the absence of glycoprotein. Coronal section through the injected telencephalic hemisphere at the level of Dp. Area outlined by dashed rectangle is enlarged. Co-expression of GFP (green) and mCherry (magenta) indicates starter cells. **(h)** Same as in (g) but with trans-complementation of zoSADG in starter neurons by co-injection of HSV1[*UAS*:zoSADG]. Left: coronal section through the injected telencephalic hemisphere. Right: coronal section through the ipsilateral OB. Expression of GFP only (green) indicates transneuronally labeled neurons. **(i)** Number of neurons in the telencephalon that expressed GFP and mCherry (putative starter neurons) or GFP alone (putative presynaptic neurons) after injection of EnvA-RVΔG and HSV1[*UAS*:TVA-mCherry] into Dp with (+) or without (-) trans-complementation with zoSADG in starter neurons (HSV1[*UAS*:zoSADG]). Left: injection into Tg[*gad1b*:Gal4] fish; right: injection into Tg[*vglut1*:Gal4] fish (right). Expression was analyzed 10 days post injection. N: number of fish. **(j)** Convergence index for the projection of transneuronally labeled neurons in the OB to Dp when EnvA-RVΔG was targeted to GABAergic neurons (viral injections into Tg[*gad1b*:Gal4] fish) or to glutamatergic neurons (injections into Tg[*vglut1*:Gal4] fish) in Dp. Expression was analyzed at 10 days post injection. N: number of fish.

To examine whether the delivery of glycoprotein to starter cells can drive transneuronal spread, we injected a mixture of HSV1[*UAS*:TVA-mCherry], HSV1[*UAS*:zoSADG] and EnvA-RVΔG-GFP into the cerebellum of Tg[*gad1b*:Gal4] fish. Unlike in the absence of glycoprotein, the number of mCherry-expressing Purkinje neurons and putative Golgi cells now declined over the incubation period of 10 days. The number of neurons expressing GFP, in contrast, was low initially but increased steeply between 6 and 10 days after injection (Fig. 4d). In the cerebellum, GFP was expressed in Purkinje cells and throughout the granular layer (Fig. 4c). The number and spatial distribution of GFP-expressing neurons in the granular layer was clearly different from mCherry expression in the absence of glycoprotein (Tg[*gad1b*:Gal4] fish injected with HSV1[*UAS*:TVA-mCherry]; Fig. 2b), indicating that GFP was expressed in granule cells. Moreover, GFP-positive neurons were found in the contralateral inferior olive (Fig. 4c). We therefore conclude that *trans*-complementation with zoSADG in starter cells promoted transneuronal spread of the rabies virus.

Transneuronal viral transfer for synaptic inputs to Purkinje neurons from the inferior olive was quantified in each individual by a convergence index that is defined as the number of transneuronally labeled neurons (here: GFP+/mCherry-neurons in the inferior olive) normalized to the number of starter cells (here: GFP+/mCherry+ Purkinje neurons)^29,32^. This index does not reflect the true convergence because an unknown number of starter neurons has disappeared at the time when neurons are counted. We nevertheless used this index to assess transneuronal spread because it has been established previously as a benchmark in rodents^29,32^. The convergence index was 1.00 ± 0.95 (mean ± s.d.; N = 4 fish; Fig. 4e), which is comparable to values reported in mammals^29^.

To corroborate these results we also examined transneuronal tracing from starter neurons in telencephalic area Dp. In this area, both glutamatergic principal neurons and GABAergic interneurons receive long-range input from mitral cells in the OB (Fig. 4f). In addition, Dp receives other telencephalic inputs that are not well characterized. When a mixture of HSV1[*UAS*:TVA-mCherry] and EnvA-RVΔG-GFP was injected into Tg[*gad1b*:Gal4] fish (Fig. 4g), GFP expression was largely restricted to a cluster of neurons that co-expressed mCherry (Fig. 4g). This cluster was located near a prominent furrow that separates an anterior from a posterior compartment of Dp, consistent with the known location of GABAergic neurons in Dp^18^. When HSV1[*UAS*:zoSADG] was added to the injected virus mixture, mCherry expression became rare while a large number of neurons in different telencephalic areas expressed GFP (Fig. 4h, i). Moreover, GFP was expressed in the olfactory bulb by neurons with the characteristic morphology of mitral cells, but not by other cells (Fig. 4h).

Similar observations were made when the infection was targeted to glutamatergic starter neurons by injecting the cocktail of viruses into Dp of Tg[*vglut1*:Gal4] fish (Fig. S9). In the absence of HSV1[*UAS*:zoSADG], GFP was co-expressed with mCherry and restricted to the injection site, while in the presence of HSV1[*UAS*:zoSADG], expression of mCherry disappeared and GFP expression became more widespread, including neurons in the OB (Fig. S9 ). Note that *vglut1* is expressed in excitatory neurons in the telencephalon but not in the OB (Fig. S10), implying that labeling of OB neurons cannot be explained by direct infection. Convergence indices for projections of mitral cells to GABAergic or glutamatergic starter neurons in Dp were 0.70 ± 0.21 and 1.31 ± 1.20 (mean ± s.d.; N = 5 and N = 4 fish, respectively; Fig. 4j). These results confirm that EnvA-RVΔG selectively infects TVA-expressing target neurons and undergoes monosynaptic retrograde transneuronal transfer when complemented with zoSADG.

To examine transneuronal viral spread at early developmental stages we injected EnvA-RVΔG-GFP into the spinal cord of Tg[*gad1b*:Gal4; *UAS*:TVA-mCherry] fish at 10 dpf and analyzed fluorescence after 6 – 7 days. In the absence of zoSADG, GFP expression was restricted to mCherry-positive neurons around the injection site (Fig. 5a). When zoSADG was supplied in *trans* by co-injecting EnvA-RVΔG-GFP with HSV1[*UAS*:zoSADG], GFP expression was observed throughout the spinal cord and in the brainstem (Fig. 5b). GFP-positive neurons distant from the injection site did usually not co-express mCherry. We therefore conclude that glycoprotein-dependent retrograde transneuronal transfer also occurs at larval stages.

**Fig 5.**
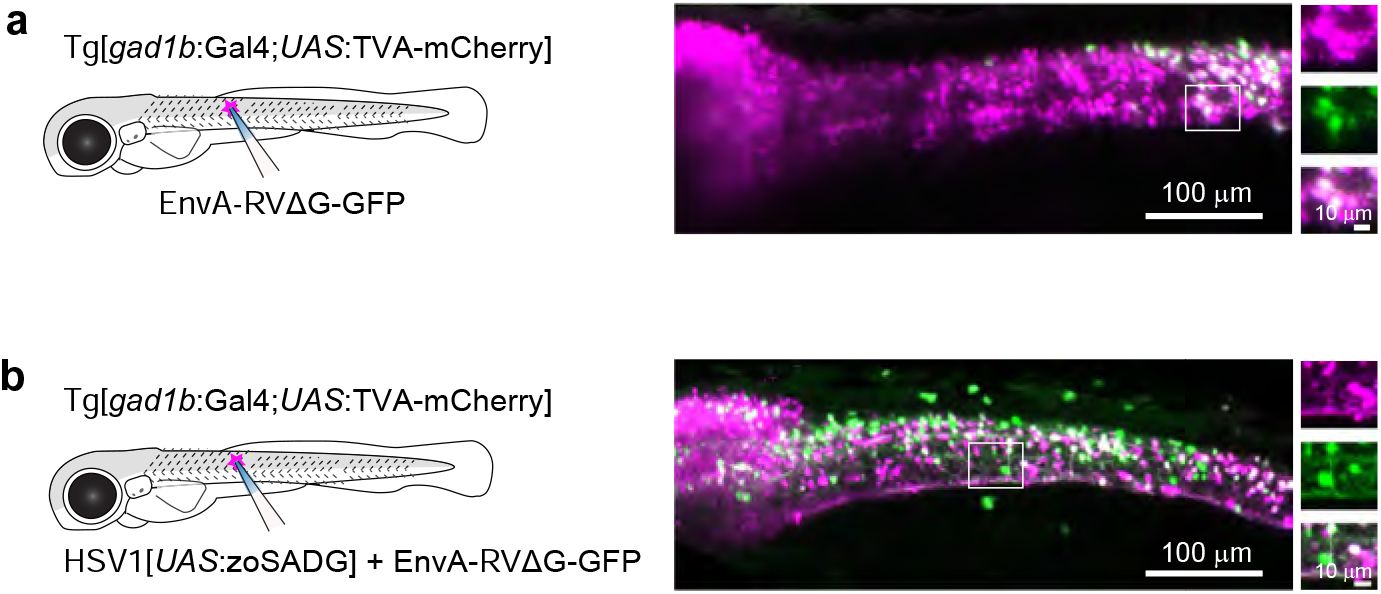
Transneuronal tracing using pseudotyped rabies virus in zebrafish larvae. **(a)** Expression of GFP (green) and TVA-mCherry (red) 6 days after injection of EnvA-RVΔG-GFP into the spinal cord of Tg[*gad1b*:Gal4;*UAS*:TVA-mCherry] fish at 7 dpf. Boxed region is enlarged on the right. **(b)** Same after co-injection of EnvA-RVΔG-GFP and HSV1[*UAS*:zoSADG] into the spinal cord of Tg[*gad1b*:Gal4;*UAS*:TVA-mCherry] fish at 7 dpf.

## Discussion

In summary, we found that viral infection and transgene expression in zebrafish can be significantly enhanced by adjusting the temperature to that of the natural viral host. HSV1 vectors can be combined with the Gal4/UAS system for intersectional gene expression strategies without obvious leakiness. We applied HSV1s to target the expression of multiple transgenes to different brain areas and neuron types in the adult zebrafish brain and in larvae. Hence, HSV1-mediated gene transfer and retrograde tracing has a wide range of potential of applications in zebrafish.

We have generated a collection of HSV1s for the direct or conditional expression of different transgenes including fluorescent markers, calcium indicators, optogenetic probes, and toxins (Supplementary Table1). This toolbox opens novel opportunities for the fast and flexible interrogation of neurons in zebrafish. Intersectional strategies can be used to target narrowly defined types of neurons by combining the genetic specificity of promoters and the Gal4/UAS system with the spatial and temporal precision of injections. For example, anatomically defined sets of projection neurons can be targeted by retrograde HSV1-mediated delivery of Gal4 and subsequent HSV1-mediated delivery of a UAS responder. Importantly, expression can be directed to specific neurons by injecting HSV1 with UAS-dependent inserts into existing Gal4 driver lines. Such approaches can immediately capitalize on the broad spectrum of Gal4 driver lines that has been created using defined promoters and random insertion strategies^33–35^. Conceivably, viral delivery of UAS responders may also be useful to overcome the slow loss of expression observed in some Gal4/UAS transgenic lines over generations because it bypasses epigenetic silencing of the highly repetitive UAS elements^36,37^.

We further established procedures for retrograde monosynaptic tracing using rabies viruses that greatly enhanced the efficiency of transneuronal transfer compared to a previous study in zebrafish^26^. Temperature adjustments were important but unlikely to be the only factor underlying the enhanced transneuronal spread because the temperature difference to the previous study was only 1.5 °C^26^. We therefore assume that increased glycoprotein expression by codon optimization, viral gene delivery, and possibly small differences in the experimental schedule also contributed significantly. Transcriptomics and measurements of odor-evoked activity revealed no signs of toxicity of RVΔG alone, indicating that transneuronally labeled neurons are healthy even though starter neurons disappear.

Transneuronal spread of rabies viruses was glycoprotein-dependent and labeled substantial numbers of putative presynaptic neurons from different starter neurons, both in adult zebrafish and in larvae. Convergence ratios were similar to those observed in rodents despite the fact that zebrafish neurons are smaller and probably receive fewer synaptic inputs. These results suggest that the efficiency of transneuronal tracing was in the same range as in rodents although quantitative comparisons are difficult. In the future, further improvements may be achieved by modifications that are currently explored in rodents such as the design of optimized glycoproteins^29^ or trans-complementation with a second viral gene^38^. The apparent long-term toxicity of the glycoprotein may be circumvented by transient expression using a negative feedback system^39^. Hence, rabies-based transneuronal tracers are promising tools for the structural and functional dissection of neuronal circuitry in zebrafish.

Retrograde tracing of connected neuronal cohorts using rabies viruses has become an important approach to decipher the functional logic of neuronal connectivity in rodents. Our results pave the way for applications of this approach in zebrafish. The small size of zebrafish facilitates the combination of neuronal circuit tracing with other methods such high-resolution imaging of neuronal activity patterns throughout large brain areas. Transneuronal tracing in the anterograde direction from starter neurons that were not genetically defined has previously been achieved using VSVs in larval zebrafish^23,24^. Recently, it has been reported that the cytotoxicity of VSVs used in initial studies can be reduced^25^, suggesting that anterograde transneuronal tracing using VSVs may complement retrograde transneuronal tracing by rabies viruses. Our experiments were designed primarily to explore HSV1 and rabies viruses as tools for structural and functional analyses of neuronal circuits and behavior. Moreover, the ability of HSV1 and rabies viruses, as well as VSVs, to infect larval neurons may be exploited for studies of development, although the temporal resolution will be limited by the time required for viral gene expression (days). Viral tools and their combination with transgenic lines and optical approaches therefore offer a wide range of experimental opportunities that have not been available in zebrafish so far.

## Acknowledgements

We thank Hans-Rudolf Hotz for offering the QuasR tools in the FMI Galaxy, Hubertus Kohler for FACS experiments, and Sébastien Smallwood and Stéphane Thiry for RNA sequencing. We thank Aya Takeoka for comments on the manuscript and the Friedrich group for stimulating discussions. This work was supported by the Novartis Research Foundation, by fellowships from the European Union (Marie Curie) and JSPS (C.S.), by the European Research Council (ERC) under the European Union’s Horizon 2020 research and innovation program (grant agreement no. 742576), and by the Swiss National Science Foundation (grant no. 31003A_172925/1).

## Author contributions

C.S. developed the methodology, designed and performed experiments, analyzed data and wrote the manuscript. R.L.N. produced HSV1. H.K.O. produced rabies virus. P. Z. performed and analyzed optogenetic and calcium imaging experiments. K.H. developed head-fix preparation and in vivo imaging technique. E.A.B. helped molecular biology to establish HSV1[*UAS*:GFP], M.T. helped to establish rabies virus tracing in zebrafish. S.H. created transgenic fish. G.B.K. supervised the project. R.W.F. supervised the project and wrote the manuscript.

## Competing interests

R.L.N. is the head of the Gene Delivery Technology Core at Massachusetts General Hospital where HSV1 can be obtained commercially.

## Methods

### Animals

Experiments were performed in adult (5 – 15 month old) zebrafish (*Danio rerio*) of both sexes. Fish were bred under standard laboratory conditions (26 – 27°C, 13 h/11 h light/dark cycle). All experiments were approved by the Veterinary Department of the Canton Basel-Stadt (Switzerland).

The following transgenic fish lines were used: Tg[*gad1b*:GFP]^33^, Tg[*gad1b*:Gal4]^18^, Tg [*vglut2a*:loxP-DsRed-loxP-GFP]^33^, Tg[*th*:Gal4]^40^, Tg[*vglut1*:GFP] (created in this study), Tg[*vglut1*:Gal4] (created in this study), Tg[*UAS*:tdTomato-CAAX]^41^ and Tg[*UAS*:TVA-mCherry] (created in this study). Note that Tg[*gad1b*:GFP] and Tg[*gad1b*:Gal4] were created using the same BAC (zC24M22).

Optogenetic experiments and odor discrimination training were performed in fish with low pigmentation that were derived by selection from a wt population. We refer to this genetic background as “Basel-golden”. These fish facilitate non-invasive optical access to the brain in adults and show no obvious impairments or behavioral alterations.

### Transgenic fish, DNA constructs and virus production

Tg[*UAS*:TVA-mCherry] fish were created using standard procedures based on the Tol2 transposon^42^. TVA-mCherry was amplified by PCR from a plasmid (gift from Dr. Uchida, Addgene plasmid #38044 ; http://n2t.net/addgene:38044 ; RRID:Addgene_38044, ref^43^) and inserted into a 5xUAS vector^34^.

Tg[*vglut1*:GFP] fish were established using the CRISPR/Cas9 method^44^. Insertion of a construct containing the hsp70 promoter was targeted at a site upstream of the *vglut1* gene using the target sequence gagagagactcgggcgcgcg. The same procedure and target sequence was used to generate Tg[*vglut1:*loxP-mCherry-loxP-Gal4]. This line was then crossed to Tg[*hspa8*:Cre-mCherry-NLS] (ZFIN ID: ZDB-ALT-201210-1), which expresses Cre ubiquitously, to generate Tg[*vglut1:*Gal4].

For HSV1 production, constructs containing *LTCMV*:DsRed, *LTCMV*:GFP, *LTCMV*:Gal4 and *LTCMV*:jGCaMP7b were created at the Gene Delivery Technology Core of the Massachusetts General Hospital. Constructs containing *UAS*:GFP, *UAS*:Venus-CAAX, *UAS*:GCamp6f, *UAS*:tdTomato, *UAS*:TeNTGFP, *UAS*:TVA-mCherry, and *UAS*:zoSADG were created using the Gateway construct described in Fig. 2A. The transcriptional blocker (Fig. 2a) and zoSADG were synthesized with the following sequences:

#### Transcriptional blocker

caataaaatatctttattttcattacatctgtgtgttggttttttgtgtgaatcgatagtactaacatacgctctccatcaaaacaaaacgaaacaaaacaaactagcaaaataggctgtccccagtgcaagtgcaggtgccagaacatttctct

#### zoSADG

atggtgcctcaggctctgctgtttgtgcctctgctggtgttccctctgtgcttcggaaaattccctatctacacaatcccggataaactgggaccttggagccctatcgacatccaccacctgagctgtcctaacaacctggtggtggaggatgagggatgcactaacctgagcggattctcctacatggagctgaaagtgggatacatcctggctatcaaggtgaacggattcacctgtacaggagtggtgactgaggctgagacatacacaaacttcgtgggatacgtgacaaccacattcaaaaggaaacacttcagacctacacctgatgcttgtagagctgcctacaactggaagatggctggtgaccctagatatgaggagtccctgcacaacccttaccctgactacagatggctgcgtacagtgaagacaacaaaagagagcctcgttatcatcagcccttccgtggccgatctcgatccatatgacagaagcctgcactctagagtgttcccaagcggaaagtgctctggcgtggcagtgtcttccacttactgctcaaccaaccacgactacaccatctggatgccagagaacccaagactgggaatgtcttgtgacatcttcactaactctagaggaaaaagagcttctaaaggatctgagacctgcggattcgtggatgagagaggactgtacaagtctctgaagggagcttgtaaactgaagctgtgtggagtgctgggactgagactgatggacggaacctgggtcagcatgcagacaagcaacgagaccaagtggtgtcctccagacaaactggtgaacctgcacgactttagaagcgatgagattgagcaccttgtggtggaggagctggtgagaaaaagagaggagtgtctggacgctctggagagcatcatgacaacaaaaagcgtgtctttcagaagactgagccacctgagaaaactggtgcctggattcggaaaggcttatacaatcttcaacaaaacactgatggaggctgatgctcactacaagagcgtgagaacatggaacgagatcctgccttctaaaggatgcctgagagtgggaggaagatgtcacccacacgtgaacggagtgttctttaacgggatcattttgg gtcccgacggcaatgtgctcatcccggaaatgcagagcagcctgctccagcaacacatggagttgctcgagagtagtgtgatacccttagtccatccactcgcagatccttccacagtgttcaaggatggtgacgaggctgaggactttgtagaggttcatctccctgatgtgcacaaccaggtgtctggagtggatctgggactgccaaactggggaaagtacgtgctgctgtctgctggagctctgaccgctctgatgctgatcatcttcctgatgacatgttgtagaagagtgaacagatctgagcctacacagcacaacctgagaggaactggaagagaggtgagcgtgacacctcagagcggaaagatcatctctagctgggagtcacataagtctggaggtgaaactagactgt ga.

All HSV1 viruses were produced by Gene Delivery Technology Core in the Massachusetts General Hospital (https://researchcores.partners.org/mvvc/about, titer: 5 x 10^9^ iu/ml). EnvA-RVΔG-GFP (titer: 2.2 x 10^9^ iu/ml) and EnvA-RVΔG-mCherry (titer: 4.2 x 10^8^ iu/ml) were produced by the FMI viral core.

### Virus injection

Virus injections were performed as described^15^ with minor modifications. Adult fish were anesthetized in 0.03 % tricaine methanesulfonate (MS-222) and placed under a dissection microscope. MS-222 (0.01 %) was continuously delivered into the mouth through a small cannula. A small craniotomy was made over the dorsal telencephalon near the midline, over the OBs, or over the cerebellum. If multiple viruses were injected in the same region, virus suspensions were mixed. Phenol red (0.05 %) was added to the suspension to visualize the injection. Micropipettes pulled from borosilicate glass were inserted vertically through the craniotomy. The depth of injections was approximately 100 μm in the dorsal telencephalon, 50 μm in the OB, 100-150 μm in the cerebellum, 300-500 μm in Dp and 50 – 200 μm in the tectum. The injected volume was 50 nl – 100 nl. After surgery, fish were kept in standard holding tanks at 35 °C – 37 °C unless noted otherwise.

Larval fish were anesthetized in 0.03 % tricaine methanesulfonate (MS-222) and placed on an agarose plate under a dissection microscope. Micropipettes pulled from borosilicate glass were inserted vertically into the target region (spinal cord, muscle or brain) without a craniotomy. The injected volume was 250 pl – 500 pl. After retraction of the pipette, fish were kept in standard tanks in an incubator at 28.5 °C, 32 °C or 35 °C.

For transneuronal rabies tracing, all components were co-injected. To test whether the efficiency of infection and transneuronal labeling can be enhanced when HSV1[*UAS*:TVA-mCherry] and HSV1[*UAS*:zoSADG] are expressed prior to the injection of EnvA-RVΔG-GFP, we also performed experiments with two separate injections. However, the number of GFP-positive neurons was not increased when EnvA-RVΔG-GFP was injected two (n = 3 fish) or four (n = 3 fish) days after HSV1[*UAS*:TVA-mCherry] and HSV1[*UAS*:zoSADG] (not shown), consistent with observations in rodents^45,46^. We assume that this observation can be explained by at least two factors. First, rabies virus appears to remain close to the injection site for an extended period of time, possibly because it has high affinity for membranes. Second, it is difficult to precisely target the two separate injections at the same neurons.

Transneuronal spread of the rabies virus was determined by quantitative analyses of neurons that expressed GFP but not TVA-mCherry. In theory, some of these neurons could be two (or more) synapses away from the starter cell if they target other neurons that received only the glycoprotein but not TVA-mCherry, if these neurons in turn were presynaptic to a starter neuron. However, because labeling was sparse and because the number of neurons receiving G only should be low, the probability of such events should be very low. Moreover, because transneuronal gene expression is observed only after a delay of >6 days, multi-step events should be very rare 10 days post infection. Multi-step events were therefore not taken into account in our quantitative analyses.

### Clearing of brain samples

We adapted the original Cubic protocol^47^ to small samples such as adult zebrafish brains. After fixation with 4 % paraformaldehyde overnight, samples were soaked with reagent 1A (10%w/v Triton, 5wt% NNNN-tetrakis (2HP) ethylenediamine and 10 %w/v urea) for 2.5 h at room temperature and for 6 h at 37 °C with mild shaking and multiple solution exchanges. Subsequently, samples were washed in PBS overnight at room temperature. On the next day, samples were treated with reagent 2 (25 % w/v urea, 50 % w/v sucrose and 10 % w/v triethanolamine) for refractive index matching, mounted in glass bottom dishes, and covered by 16 x 16 mm cover glasses to avoid drift. Images were acquired using a Zeiss 10x water-immersion objective lens (N.A. = 0.45) on an upright Zeiss LSM 700 confocal microscope.

### Immunohistochemistry

For rabies virus tracing in the cerebellar circuit, GFP and mCherry signals were detected by immunocytochemistry. Brain samples were fixed overnight in 4 % paraformaldehyde and sectioned (100 μm) on a Leica VT1000 vibratome. Primary antibodies were anti-GFP (Thermofisher, A10262, 1:200) and anti-RFP (5F8, chromotek, 1:200). Secondary antibodies were conjugated to Alexa Fluor 488 or 594 (Invitrogen, 1:200).

### Neuron counts

For whole brain imaging (Fig. 1c, 1g and S1d), z stacks (7-10 μm steps) were acquired using an upright Zeiss LSM 700 confocal microscope with a 10x objective (water, N.A. = 0.45, pixel size 1.25 μm). GFP or DsRed-expressing neurons in the dorsal telencephalon were counted manually.

Z stacks from individual brain slices (1 – 2 um steps, Fig. 2f, 4d, 4i, 4j, and S4c) were acquired using an upright Zeiss LSM 700 confocal microscope with a 20x objective (air, N.A. = 0.8, pixel size 0.625 um). For non-rabies injections, a single slice, containing the largest number of labelled neurons, was chosen from each brain sample for cell counting. For rabies injections, all slices were used for cell counting. Cells expressing GFP and/or mCherry in specific regions (cerebellum, telencephalon, olfactory bulb) were counted manually.

### Optogenetics and imaging *in vivo*

Multiphoton calcium imaging and simultaneous ChrimsonR stimulation was performed as described^48^ using a modified B-scope (Thorlabs) with a 12 kHz resonant scanner (Cambridge Technology) at excitation wavelengths of 930 nm (GCaMP6f) or 1100 nm (tdTomato) and an average power under the objective of 30 mW at 930 nm. Optogenetic stimulation with ChrimsonR was performed using an LED (UHP-T-595, Prizmatix; 595 nm). Light paths for imaging and stimulation were combined using a dichroic mirror (ZT775sp-2p, Chroma). Emitted light was split using a second dichroic mirror (F38-555SG, Semrock), band-pass filtered with a 525/50 filter (Semrock) for GCaMP6f imaging and with a 607/70 filter (Semrock) for tdTomato, and focused onto a GaAsP photomultiplier (H7422, Hamamatsu). The signal was amplified (DHPCA-100, Femto), digitized at 800 MHz (NI5772, National Instruments), and band-pass filtered around 80 MHz using a digital Fourier-transform filter implemented on an FPGA (NI5772, National Instruments). LED activation was synchronized to the turnaround phase of the resonant scanner when no data were acquired. Images were acquired at 60 Hz with a resolution of 750 x 400 pixels, corresponding to a field of view of 300 µm x 250 µm. Images were acquired sequentially in four different focal planes by moving the objective (Nikon 16x, 0.8 NA) with a piezo-electric linear actuator (Physik Instrumente; effective frame rate: 15 Hz per plane). Anatomical snapshots were generated by averaging 1000 images in the absence of optogenetic stimulation.

Basel-golden fish were head-fixed in a custom chamber and neurons in the dorsal telencephalon were imaged through the intact skull^5^. Optogenetic stimuli (500 ms duration) were applied randomly every 6 – 11 s. The average power of each stimulus was chosen at random to be 1.3 mW, 2.4 mW, 4.7 mW, or 8.6 mW under the objective. The total duration of an imaging session was 11 min.

Raw images were full frame registered to correct for motion. Regions of interest were manually selected based on neuronal co-expression of GCaMP6f and ChrimsonR-tdTomato (N = 11 neurons). Raw fluorescence traces were calculated as mean of the pixel values in a given region of interest in each imaging frame. Raw traces were then corrected for slow drift in fluorescence using an 8th-percentile filtering with a 15 s window^49^. ΔF/F traces were computed by dividing raw fluorescence trace by the median calculated over the entire fluorescence distribution for each region of interest. Responses were pooled across neurons and pulses of the same intensity, and the resulting population responses were normalized by subtracting average population activity in a 1 s baseline window prior to the pulse. The standard error of the mean population response was computed over average responses of individual neurons.

### Calcium imaging in the OB

Calcium imaging in the adult OB was performed 4 – 10 days after viral injections in an explant preparation of the entire brain and nose^18,50^ that was continuously superfused with artificial cerebrospinal fluid (ACSF) containing (in mM) 124 NaCl, 2 KCl, 1.25KH2PO4, 1.6 MgSO4, 22 D-(+)-Glucose, 2 CaCl2, 24 NaHCO3, pH 7.2^51^. Oregon Green 488 1,2-bis-(*o*-aminophenoxy)-ethane-*N*,*N*,*N*,*N*-tetraacetic acid, tetraacetoxymethyl ester (OGB-1; Thermo Fisher Scientific) was injected into the OB as described^18^. Two-photon calcium imaging started >1 hr after dye injection. Odors were prepared and delivered to the nose for 5 s as described^52^. Inter-stimulus intervals (ISIs) were >2 min.

Multiphoton calcium imaging was performed using a custom-built microscope with a 20x water immersion objective (NA 1.0; Zeiss) and galvo scanners^50^. Excitation wavelengths were 930 nm (OGB-1) or 1010 nm (mCherry). The average power under the objective was 50 mW at 930 nm and 20 mW at 1010 nm. The emitted light was split by a dichroic mirror (DMSP550L, Thorlabs), band-pass filtered with a 515/30 filter (Chroma) or with a 641/75 filter (Semrock), and collected with a GaAsP photomultiplier (H7422-40MOD or H11706P-40, Hamamatsu). Images were acquired at 8 Hz using Scanimage 5.5-1 (Vidrio Technologies, LLC)^53^ with a resolution of 256 x 256 pixels.

### Tissue dissociation and cell sorting

EnvA-RVΔG-GFP was injected into one or both OBs of Tg[*gad1b*:Gal4;*UAS*:TVA-mCherry] fish. Cells were dissociated and sorted as per previous protocol^54^ with modification for fish. Briefly, after 3 – 4 days at 36 °C, fish were anesthetized by cooling to 4 °C and decapitated in ACSF supplemented with 50 mM 2-Amino-5-phosphonovaleric acid (APV), 20 mM 6,7-Dinitroquinoxaline-2,3-dione (DNQX), 5 mg/ml actinomycin D (ACT-D), tetrodotoxin (TTX) 100 nM and 10 g/l Trehalose. OB samples were taken and harvested and kept in ACSF. After pooling OB samples from 3 – 5 fish, samples were treated with pronase-mix (1 g/l protease type xiv and 33 mg/l of collagenase in ACSF) for 10 minutes and the solution was replaced with fresh ACSF containing 1 % FBS. Samples were then triturated gently with small custom-made glass pipettes and stored on ice. DAPI was added to the samples to detect dead cells and cells were sorted based on GFP and mCherry fluorescence using a 70 or 100 μm nozzle (BD FACSAria III; BD Biosciences). Sorted cells were kept in a lysate buffer and stored at -80 °C until further processing.

### RNA sequencing

RNA was purified using a single-cell RNA purification Kit (Norgen). mRNA-seq libraries were generated using the SmartSeq2 approach^55^ with the following modifications: For cDNA pre-amplification, up to 10 ng of RNA was used as input and reverse transcription was performed using Superscript IV (Thermo Fisher Scientific - 50 °C for 10 min, 80 °C for 10 min). Amplified cDNA (1 ng) was converted to indexed sequencing libraries by tagmentation, using in-house purified Tn5^56^ and Illumina Nextera primers. Libraries were sequenced on an Illumina HiSeq2500, as 50 bp single-end reads.

Sequenced reads were pre-processed with preprocessReads from the Bioconductor package QuasR (version 1.24.2) ^57^)with default parameters except for Rpattern = “CTGTCTCTTATACACATCT”. Processed reads were then aligned against the chromosome sequences of fish genome (danRer11) using qAlign (QuasR) with default parameters except for aligner = “Rhisat2”, splicedAlignment = “TRUE”, and auxiliaryFile = “mCherry_GFP1.fa”. mCherry and GFP sequences matched the sequences in the plasmids used to generate transgenic fish (Tg[*UAS*:TVA-mCherry]) or rabies virus^58^.

Raw gene counts were obtained using qCount (QuasR) with default parameters and ZFIN gene models (https://zfin.org/downloads/zfin_genes.gff3, downloaded 18-Jul-2019) or the auxiliary file as query. The count table was filtered to remove genes which had less than 2 samples with at least 1 cpm. Differential gene expression was calculated with the Bioconductor package edgeR (version 3.26.4)^59^ using the quasi-likelihood F-test after applying the calcNormFactors function, obtaining the dispersion estimates and fitting the negative binomial generalized linear models. The following threshold was applied: Significant differences in gene expression were detected by applying a threshold of abs(logFC (fold change)) >3, logCPM (counts per million reads mapped to the annotation) >3, and FDR (False Discovery Rate)<0.05. Stress marker genes (GO:0033554) and cell death marker genes (GO:0008219) were chosen from Gene Ontology database (AmiGo2: http://amigo.geneontology.org/amigo/landing). Gene ontology term for differentially expressed genes were found using GENERIC GENE ONTOLOGY (GO) TERM FINDER (https://go.princeton.edu/cgi-bin/GOTermFinder).

### Odor discrimination training and analysis of behavior

In the experimental group, Basel-golden fish were injected with HSV1[*LTCMV*:jGCaMP7b] into both OBs and kept at 36 °C for 2 days prior to behavioral training. Associative conditioning was performed as described^6,18^. Briefly, individual fish were acclimated to the behavioral setup without food for 1 – 3 days and subsequently trained to associate one odor stimulus (CS^+^: alanine) with a food reward, whereas a second odor stimulus (CS^−^: tryptophan) was not rewarded. Each odor was infused into the tank for 30 s nine times per day in an alternating sequence (inter-trial interval: 20 min) for five consecutive days. A small amount of food was delivered at a specific location at the end of the presentation of the CS^+^ but not the CS^-^. 3D swimming trajectories were reconstructed from videos acquired by two orthogonal cameras (Logitech HD Pro C920). The following behavioral components were extracted from the trajectories: swimming speed, elevation in water column, presence in the reward zone, surface sampling, distance to the odor inflow and rhythmic circular swimming. The components were combined into a compound score of appetitive behavior as described^6^. The learning index was calculated as the difference between the mean behavioral scores in response to the CS^+^ and CS^−^ during the final day of training (last nine trials with each odor) in each fish.

## Data availability

Transcriptomic data generated in this study have been deposited at ArrayExpress (accession number: E-MTAB-11083).

Requests for HSV1 and the vectors shown in Fig. 2a should be addressed to R.L.N.

The data that support the findings of this study are available from the corresponding authors upon reasonable request.

## Code availability

All code used in this study is available from the corresponding authors upon reasonable request.

## Supplementary Figure Legends

**Fig S1.**
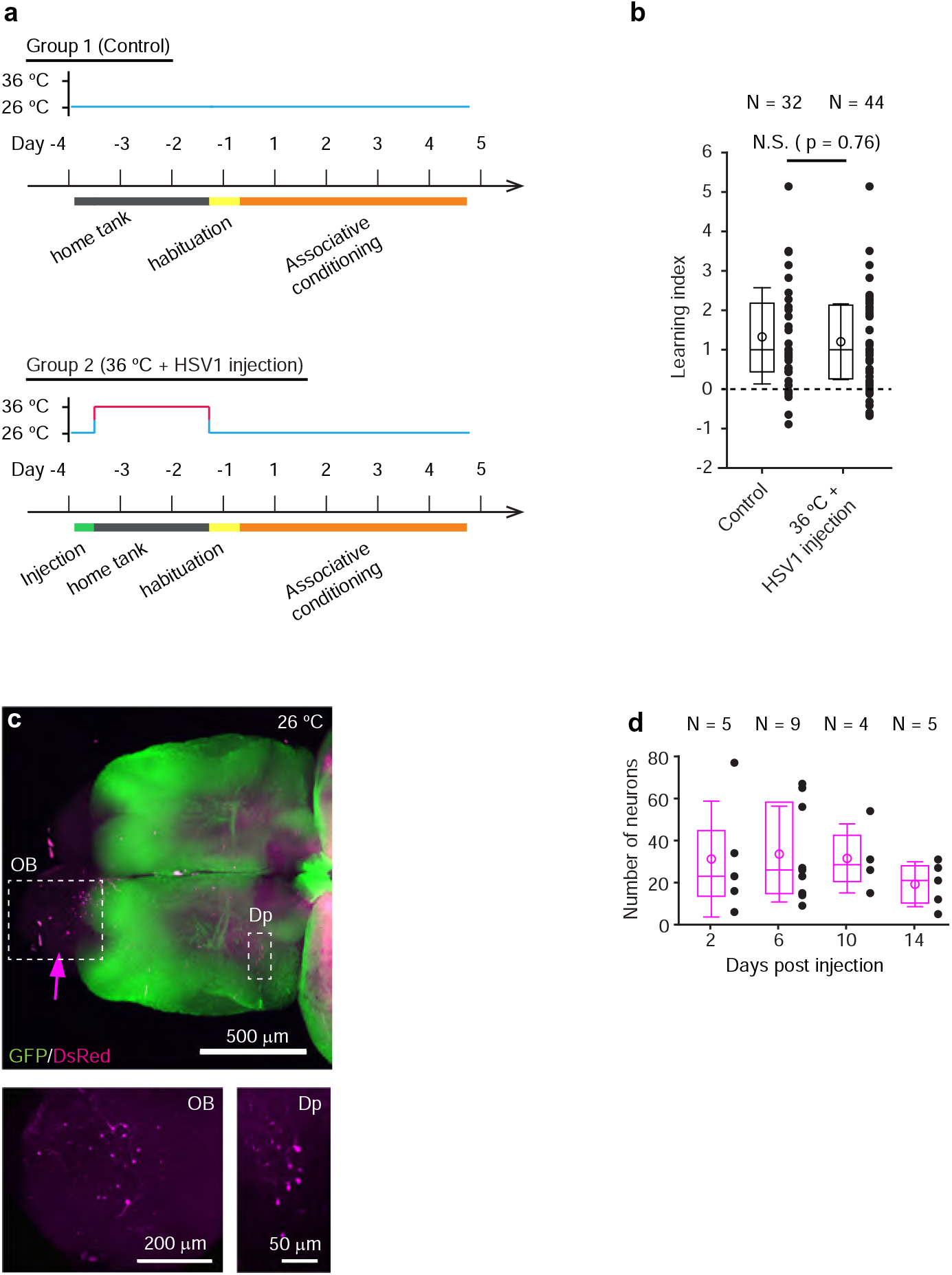
Temperature-dependence and time course of HSV1-mediated gene expression. **(a)** Sequence of virus injections, temperature changes and associative conditioning to assess effects of experimental manipulations on discrimination learning. **(b)** Learning index (behavioral discrimination score) on the last day of training. Plot symbols represent data from individual fish; box plots show median and 25^th^ and 75^th^ percentiles, circles and error bars indicate mean and s.d. over individual fish. N: number of fish. Control fish (group 1) were not injected and kept at standard laboratory temperature. The experimental group (group 2) was injected with HSV1[*LTCMV*:jGCaMP7b], an amplicon type of HSV1 with an insert encoding the calcium indicator GCamp7b^19^ under the control of a non-specific promoter for long-term expression (*LTCMV*), and subsequently kept at 36 °C for two days before training **(a)**. Performance was not significantly different between groups (p = 0.76, Wilcoxon rank sum test). **(c)** Expression of DsRed (magenta) six days after injection of HSV1[*LTCMV*:DsRed] into the OB (arrow) of an adult Tg[*vglut1*:GFP] fish kept at 26 °C (maximum projection of confocal stack). Boxed areas (OB and Dp) are enlarged below. The number of DsRed-expressing neurons is low compared to DsRed expression at 36 °C (Fig. 1). **(d)** DsRed expression in the dorsal telencephalon at different timepoints after injection of HSV1[*LTCMV*:DsRed] into the ipsilateral OB. Fish were kept at 36 °C. Black dots represent data from individual fish, box plot indicates median and 25^th^ and 75^th^ percentiles, circles and error bars indicate mean and s.d. over individual fish. N: number of fish.

**Fig S2.**
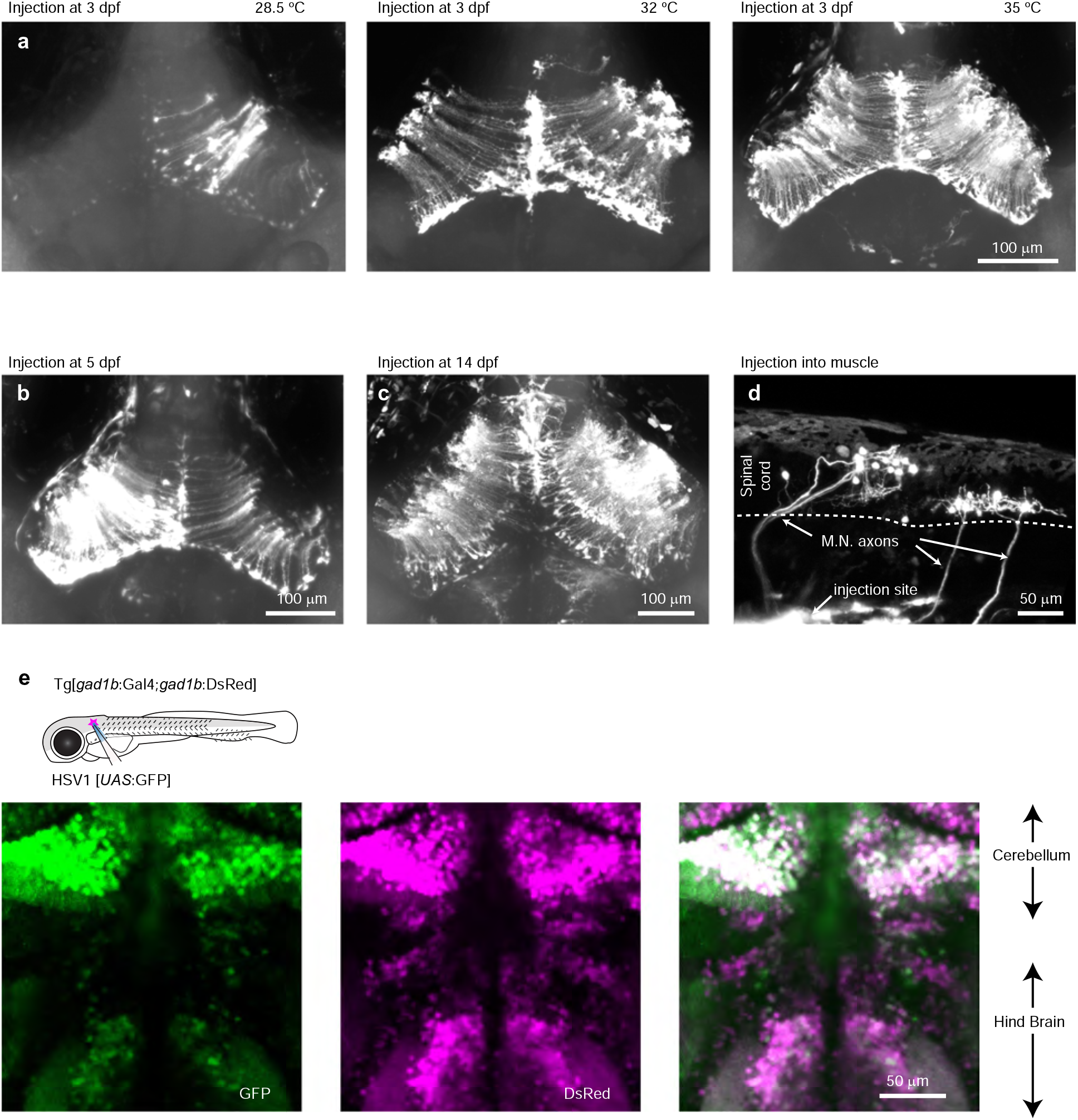
HSV1-mediated gene delivery in larvae zebrafish and Gal4/UAS. **(a)** Expression of GFP 48 h after injection of HSV1[*LTCMV*:GFP] into the optic tectum of zebrafish larvae (3 dpf; maximum intensity projection of confocal stack). Larvae were kept after the injection at 28.5, 32 or 35 °C. **(b)** Expression of GFP 48 h after injection of HSV1[*LTCMV*:GFP] into the optic tectum of a larva at 5 dpf (maximum intensity projection of confocal stack). The larva was kept after the injection at 35 °C. **(c)** Expression of GFP 48 h after injection of HSV1[*LTCMV*:GFP] into the optic tectum of a larva at 14 dpf (maximum intensity projection of confocal stack). The larva was kept after the injection at 35 °C. **(d)** Expression of GFP 48 h after injection of HSV1[*LTCMV*:GFP] into trunk muscles at 7 dpf (maximum intensity projection of confocal stack). The larva was kept after the injection at 35 °C. Note retrograde labeling of motor neurons (M.N.). **(e)** Expression of GFP 48 h after injection of HSV1[*UAS*:GFP] into the hindbrain of a Tg[*gad1b*:Gal4; *gad1b*:DsRed] larva at 7 dpf (maximum intensity projection of confocal stack). The larva was kept after the injection at 35 °C. Note co-localization of DsRed and GFP in hindbrain and cerebellum.

**Fig S3.**
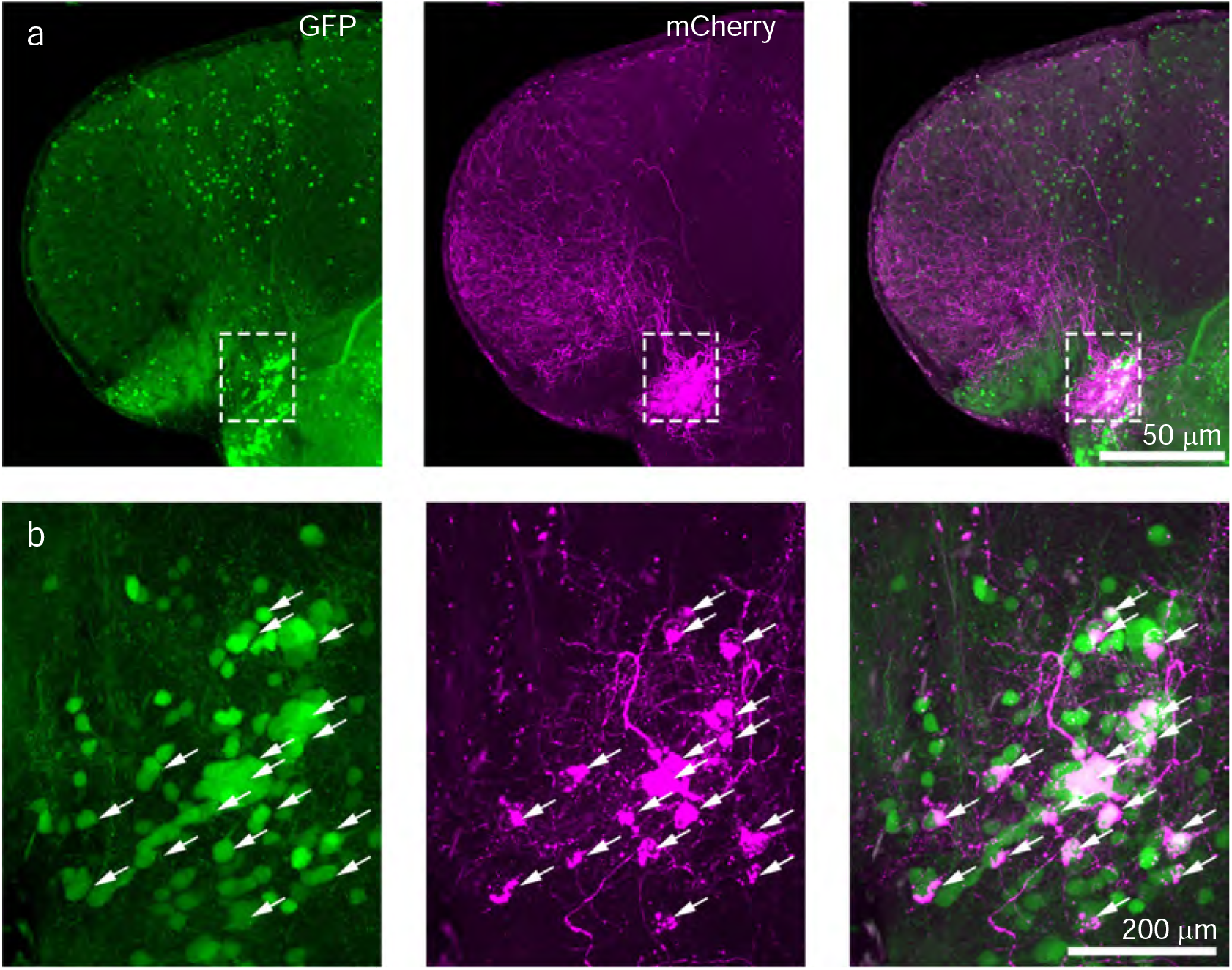
Targeting of GABAergic neurons in the telencephalon. **(a)** Coronal section through the telencephalon at the level of Dp after injection of HSV1[*UAS*:TVA-mCherry] into Tg[*gad1b*:Gal4; *gad1b*:GFP] double transgenic fish. The injection was targeted to a volume around Dp. mCherry was expressed predominantly in a cluster of GFP-positive neurons associated with Dp. Note long-range projections of mCherry-expressing neurons to multiple telencephalic areas. **(b)** Enlargements of boxed region in (a). Arrowheads indicate GFP+/mCherry+ neurons.

**Fig S4.**
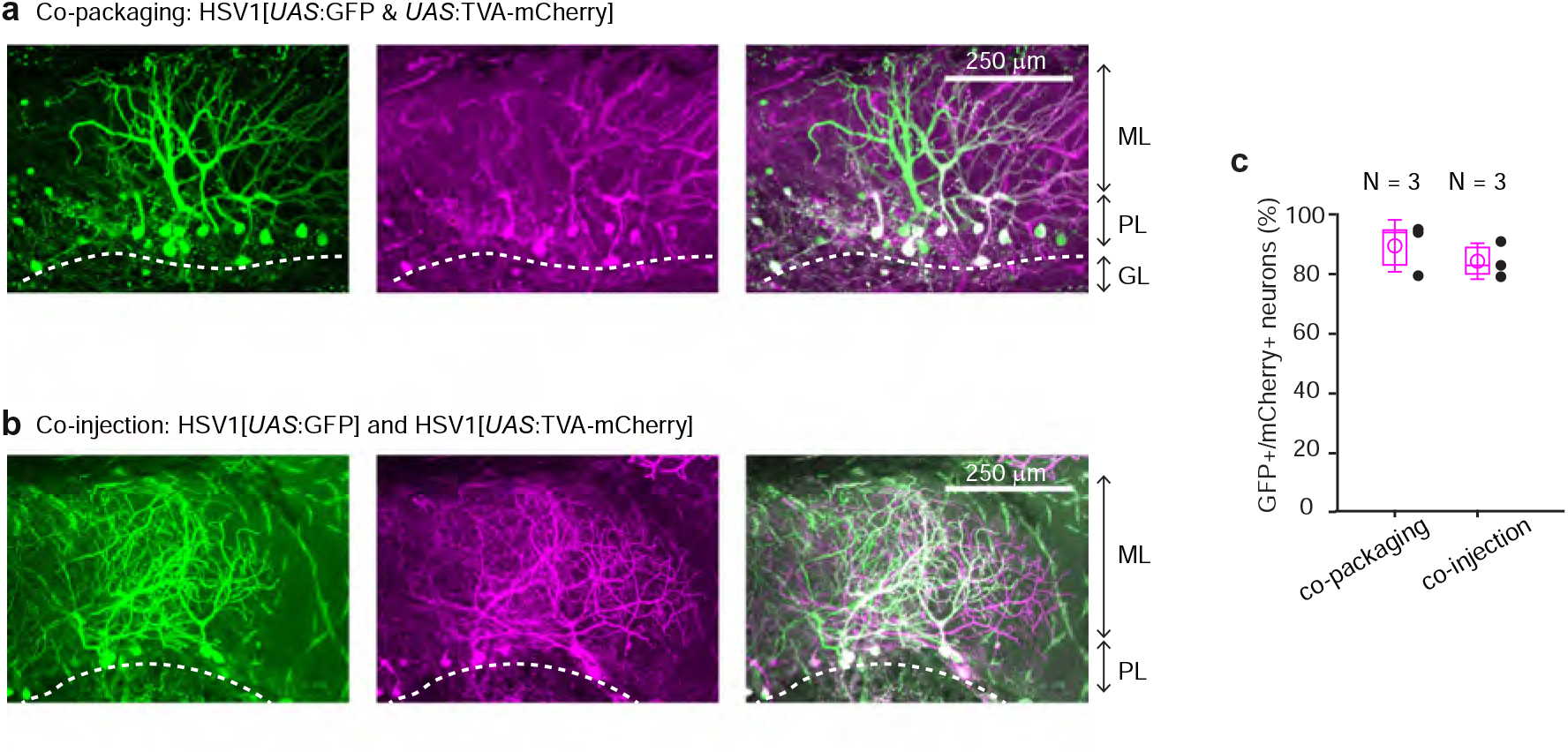
Co-packaging of 2 different viruses does not facilitate co-infection of 2 viruses. **(a)** Expression of GFP and mCherry after injection of HSV1[*UAS*:GFP & *UAS*:TVA-mCherry] into the cerebellum of Tg[*gad1b*:Gal4] fish. In this virus, two expression constructs, *UAS*:GFP and *UAS*:TVA-mCherry, are packaged into the same virus particles. Expression is observed in Purkinje neurons and in putative Golgi cells. Note high rate of co-expression of GFP and mCherry. ML: molecular layer; PL: Purkinje layer; GL: granular layer. **(b)** Expression of GFP and mCherry in the Purkinje layer after co-injection of two independent viruses (HSV1[*UAS*:GFP] and HSV1[*UAS*:TVA-mCherry]) into the cerebellum of Tg[*gad1b*:Gal4] fish. Note that the rate of co-expression was high even though GFP and mCherry were delivered by separate viruses. Note also that the overall expression was sparse, implying that co-expression was unlikely to occur by chance. **(c)** Percentage of GFP and mCherry-expressing neurons among all fluorescent neurons. Filled circles represent data from individual fish, box plot indicates median and the 25^th^ and 75^th^ percentiles, and open circles indicate mean over individual fish. N: number of fish.

**Fig S5.**
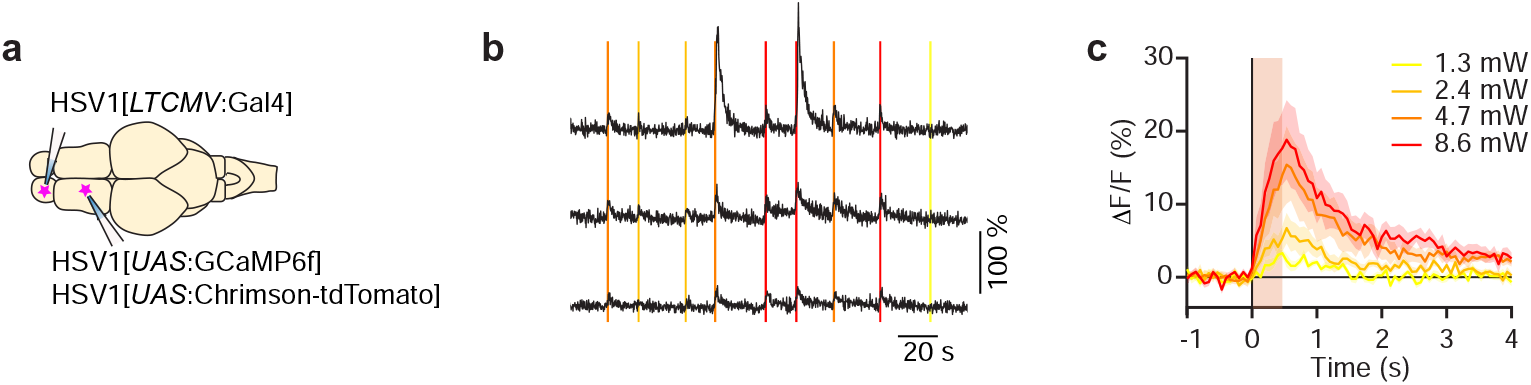
Optogenetic manipulations using HSV1. **(a)** Schematic: injection of HSV1[*LTCMV*:Gal4] into the OB and subsequent co-injection of HSV1[*UAS*:GCaMP6f] and HSV1[UAS:Chrimson-tdTomato] into the dorsal telencephalon of wt fish. **(b)** Simultaneously recorded calcium transients evoked by optical stimulation of different light intensity (vertical lines) in three example neurons. **(c)** Mean change GCaMP6f evoked by optical stimulation of different light intensity (N = 11 neurons; 11 – 18 light stimuli at each intensity). Shading shows SEM is over cell.

**Fig S6.**
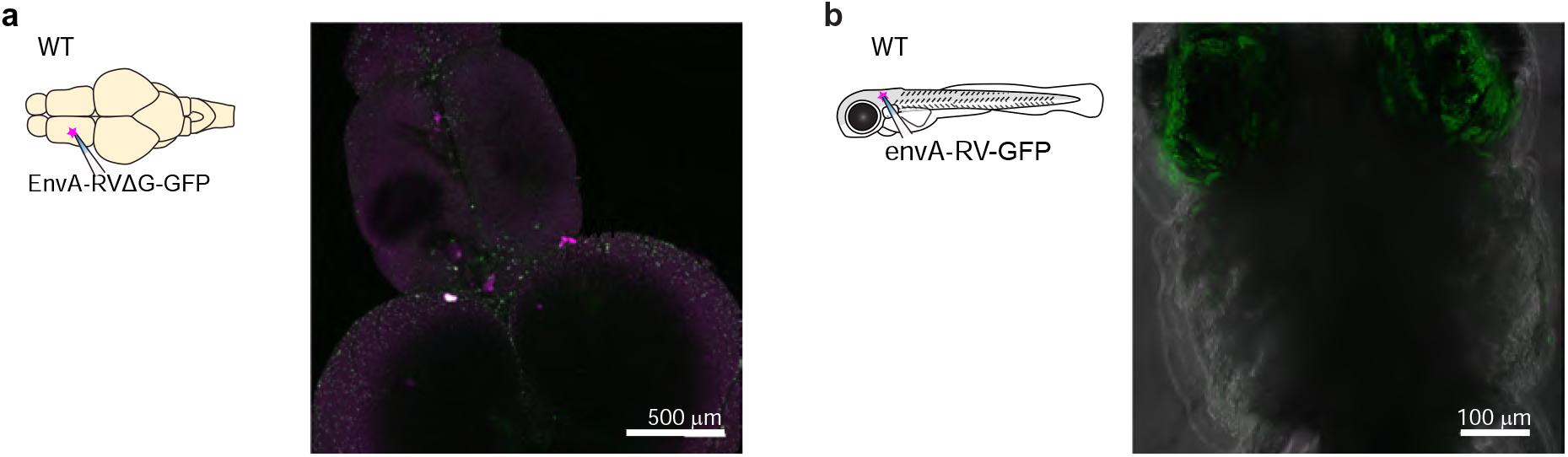
Injection of pseudotyped rabies virus does not infect neurons in the absence of TVA. **(a)** Absence of expression after injection of EnvA-RVΔG-GFP into the telencephalon of adult wt fish. **(b)** Absence of expression after injection of EnvA-RVΔG-GFP into the optic tectum of wt fish at 7 dpf.

**Fig S7.**
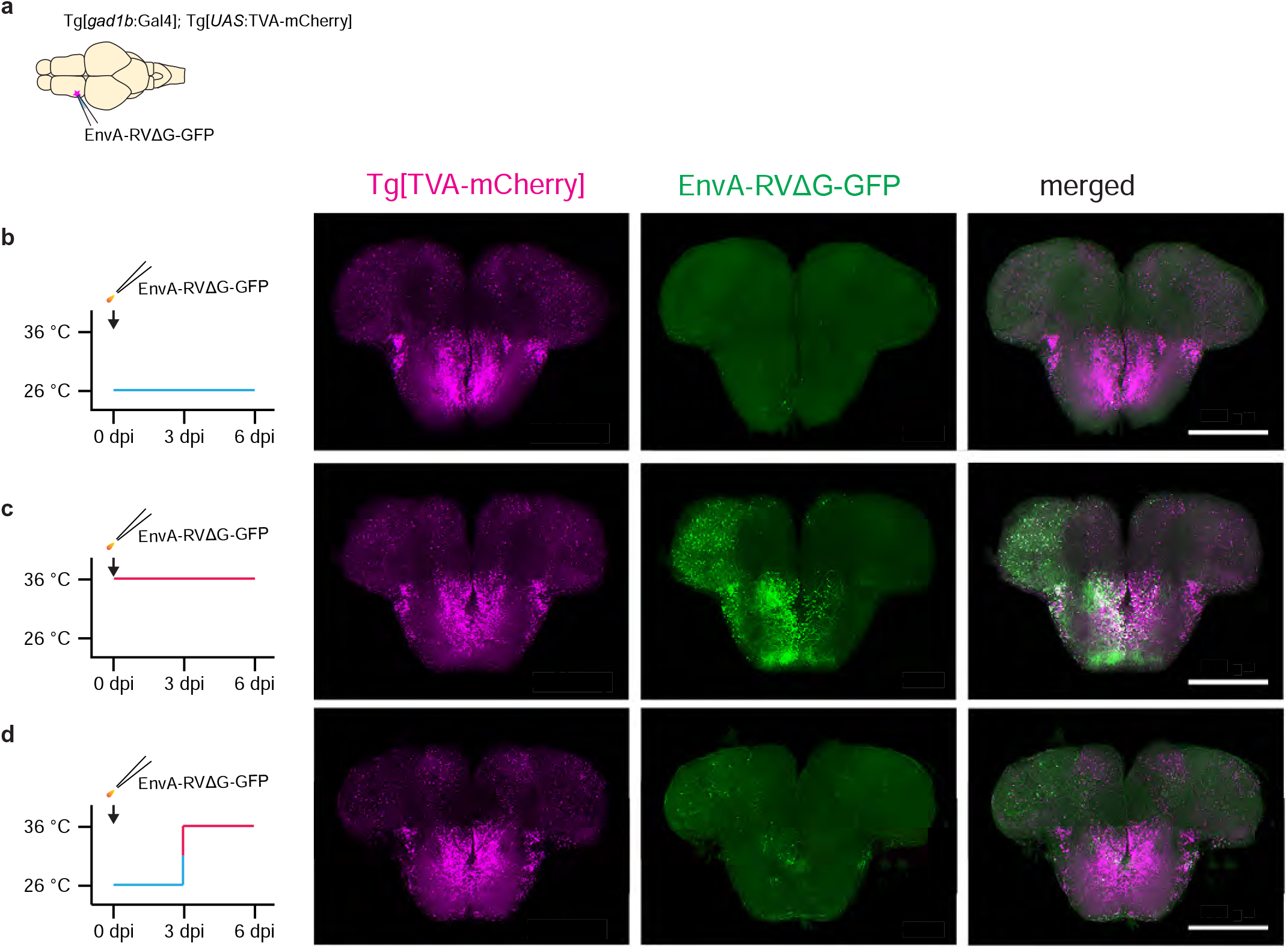
Temperature-dependence of infection by rabies virus. **(a)** Experimental scheme: Rabies virus (EnvA-RVΔG-GFP) was injected into the telencephalon of transgenic fish expressing TVA-mCherry in GABAergic neurons (Tg[*gad1b*:Gal4; *UAS*:TVA-mCherry]). **(b)** Expression of TVA-mCherry and GFP when fish were kept at 26 °C for 6 days after injection. Note almost complete absence of GFP expression. **(c)** Expression of TVA-mCherry and GFP when fish were kept at 36 °C for 6 days after injection. Note strong GFP expression. **(d)** Expression of TVA-mCherry and GFP six days after injection when the housing temperature was increased from 26 °C to 36 °C 3 days after injection. GFP expression was weak and sparse.

**Fig S8.**
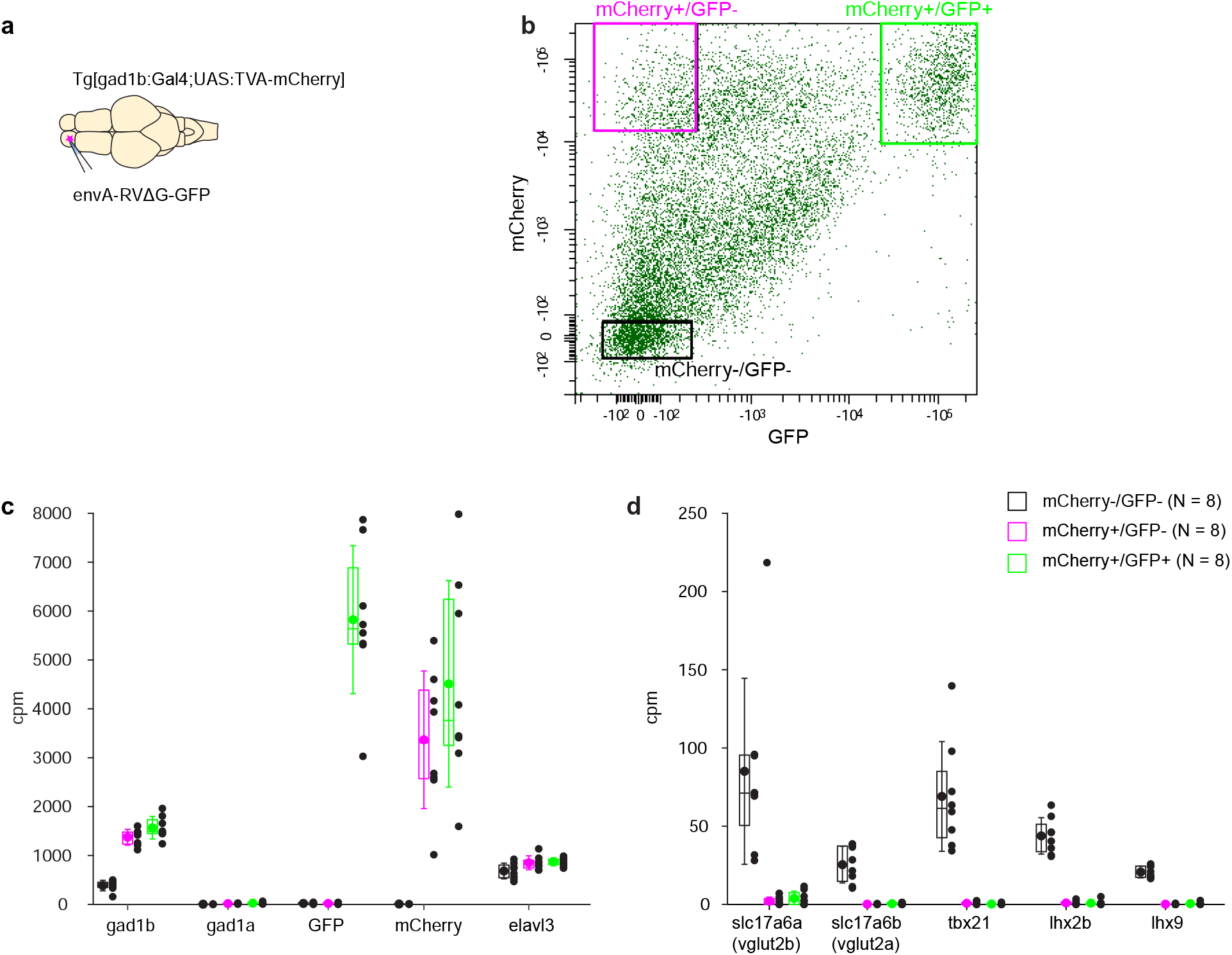
Analysis of gene expression in GABAergic neurons. **(a)** Schematic: injection of EnvA-RVΔG-GFP into the OB of adult Tg[*gad1b*:Gal4;*UAS*:TVA-mCherry] fish. **(b)** Example of FACS analysis of GFP and mCherry expression. Boxes depict cells selected as mCherry+/GFP+ (EnvA-RVΔG-GFP infected gad1b neurons), mCherry+/GFP- (non-infected gad1b neurons), mCherry-/GFP- (negative control containing other OB cells). gad1b is one of two isoforms of gad1 that are expressed differentially in GABAergic neurons. **(c)** Expression of marker genes (x-axis) in infected gad1b neurons (mCherry+/GFP+; green), non-infected gad1b neurons (mCherry+/GFP-; magenta), and other OB cells (mCherry-/GFP-; black). Cells classified as gad1b-positive by fluorescence markers, but not other cells, expressed *gad1b* but not *gad1a*, the other *gad1* isoform. Expression of fluorescent marker genes followed the detection of fluorescent markers by FACS. The neuronal marker *elav3* was present in all three pools. Plot symbols represent data from individual samples; box plots show median and 25^th^ and 75^th^ percentiles, circles and error bars indicate mean and s.d. over individual samples (N = 8 samples). **(d)** Expression of negative markers for GABAergic neurons. The selected marker genes (*slc17a6a*, *slc17a6b*, *tbx21*, *lhx2b* and *lhx9*) should be expressed in mitral cells of the OB and other excitatory neurons but not in GABAergic neurons. Consistent with this expectation, expression of all negative markers was low or absent in pools of *gad1b* cells selected by FACS (N = 8 samples).

**Fig S9.**
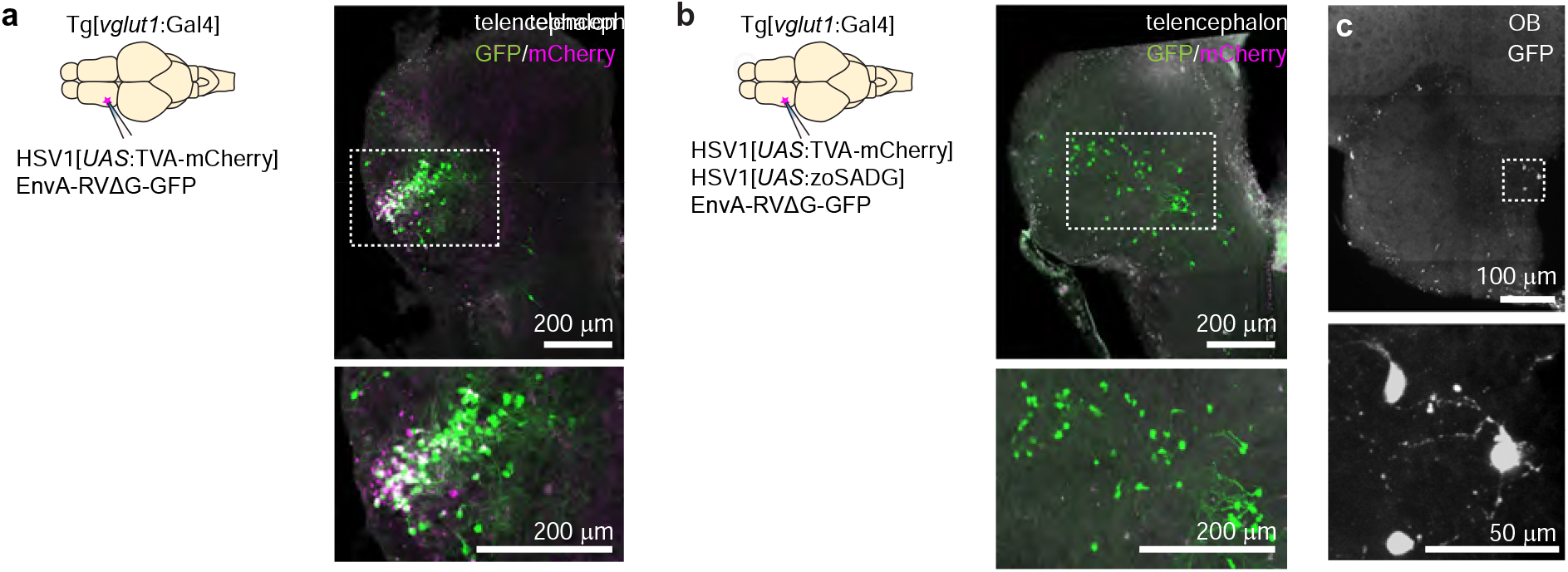
Transneuronal tracing using pseudotyped rabies virus from *vglut1*+ neurons in Dp in adult zebrafish. **(a)** Co-injection of EnvA-RVΔG-GFP and HSV1[*UAS*:TVA-mCherry] into Dp of Tg[*vglut1*:Gal4] fish in the absence of glycoprotein. Coronal section through the injected telencephalic hemisphere at the level of Dp. Area outlined by dashed rectangle is enlarged. Co-expression of GFP (green) and mCherry (magenta) indicates starter cells. **(b)** Same as in (a) but with trans-complementation of zoSADG in starter neurons by co-injection of HSV1[*UAS*:zoSADG]. Left: coronal section through the injected telencephalic hemisphere. Right: coronal section through the ipsilateral olfactory bulb. Expression of GFP only (green) indicates transneuronally labeled neurons.

**Fig S10.**
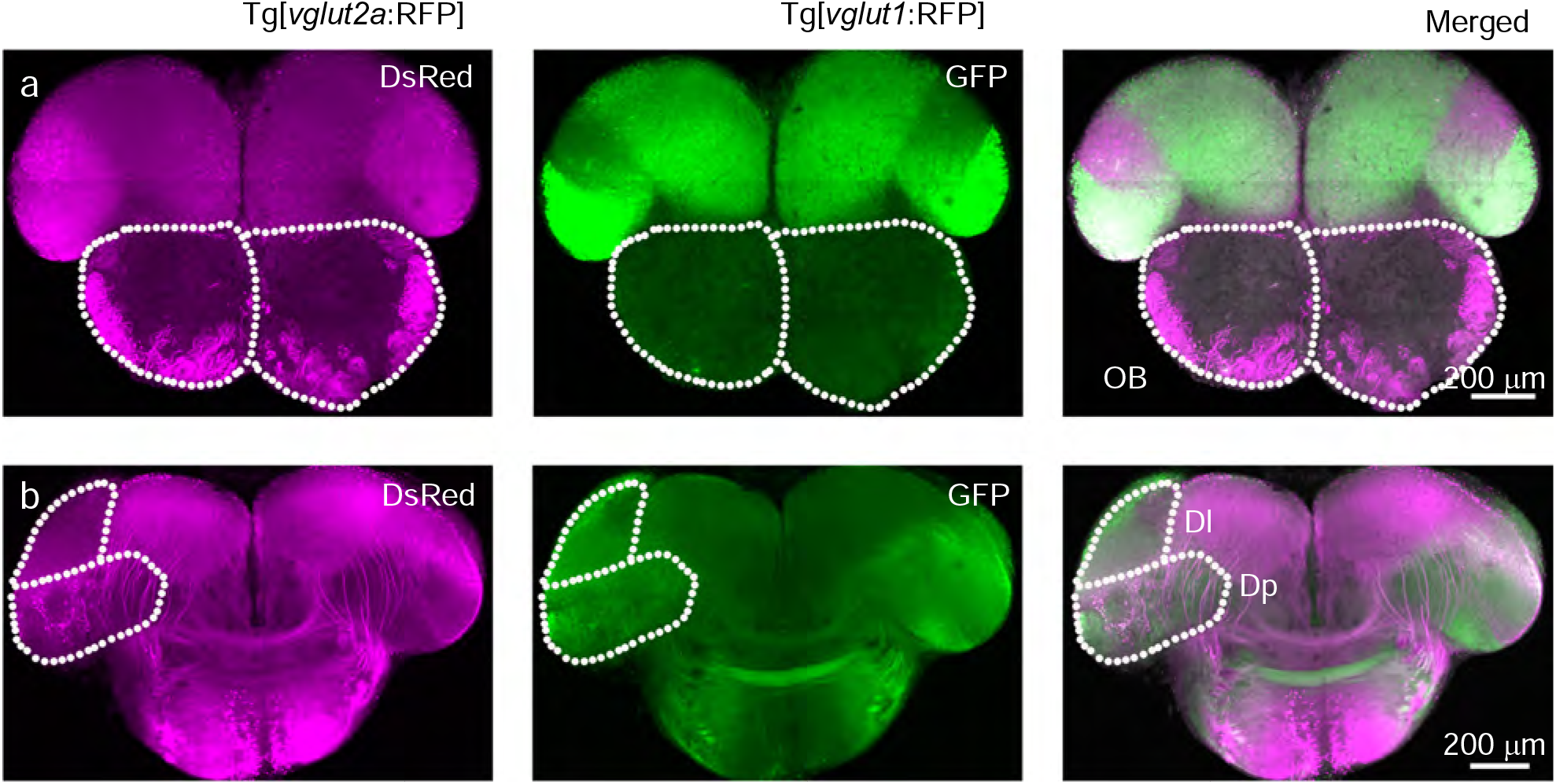
Expression pattern of *vglut1* and *vglut2* in olfactory bulb and Dp. **(a)** Coronal cross sections through the OB and anterior telencephalon from Tg[*vglut2a*:RFP; *vglut1*:GFP] double transgenic fish. Note that *vglut2a* (magenta) is expressed by axons of olfactory sensory neurons innervating glomeruli in the OB and by a subset of mitral cells, while expression of *vglut1* (green) in the OB is weak or absent. Dotted lines outline OBs. **(b)** More posterior coronal cross sections through the telencephalon of the same fish at the level of Dp. Note that expression of *vglut2a* and *vglut1* in the telencephalon are largely complementary. Neurons in Dp express primarily *vglut1*. Dotted areas indicate the dorsal lateral telencephalic area (Dl) and Dp.

**Movie1:** Swimming behavior of adult zebrafish injected with an HSV1[*LTCMV*:DsRed] into the OB at 27 °C. Fish were kept at 27 °C for 10 days after the injection.

**Movie2:**

Swimming behavior of adult zebrafish injected with an HSV1[*LTCMV*:DsRed] into the OB at 37 °C. Fish were kept at 37 °C for 10 days after the injection.

**Movie3:** Swimming behavior of zebrafish larvae injected with an HSV1[*LTCMV*:GFP] into the optic tectum at 35 °C. Fish were injected at 7 dpf and kept for 2 days at 35 °C.

**Movie4:** Effect of TeNT expression in GABAergic neurons of the cerebellum on swimming behavior. Left: Tg[*gad1b*:Gal4] fish 3 days after injection of HSV1[*UAS*:GFP] into the cerebellum (control). Right: Tg[*gad1b*:Gal4] fish 3 days after injection of HSV1[*UAS*:TeNT-GFP] into the cerebellum. Top: side view; bottom: top view of the same tanks.

**Table 1.** List of HSV1 available. Requests for HSV1 and the destination vector shown in Fig. 2a should be addressed to R.L.N.

| Promoter | Transgene | Transgene description | Comments |
| --- | --- | --- | --- |
| LTCMV | GFP | Green fluorescent protein | Fig. S2 |
|  | DsRed | Red fluorescent protein | Fig. 1b, Fig. S2c |
|  | mCherry | Red fluorescent protein |  |
|  | tdTomato | Red fluorescent protein |  |
|  | tagBFP-CAAX | Blue fluorescent protein, membrane targeted |  |
|  | Venus-CAAX | Yellow fluorescent protein (YFP), membrane targeted |  |
|  | TVA-mCherry | TVA-mCherry fusion |  |
|  | GCamp6s | Green fluorescent calcium indicator |  |
|  | Gal4 | Gal4 | Fig. 2e, Fig. S5 |
|  | ChR2-YFP | Channelrhodopsin-2-YFP fusion |  |
|  | ChR2-mCherry | ChR2-mCherry fusion |  |
|  | eNpHR3.0 | Halorhodopsin |  |
|  | H2B_GCamp6s | Green fluorescent calcium indicator, targeted to nucleus |  |
|  | jGCamp7s | Green fluorescent calcium indicator |  |
|  | jGCamp7b | Green fluorescent calcium indicator | Fig. S1 b |
| UAS | GFP | Green fluorescent protein | Fig. 2b, Fig. S2e, Fig. S4 |
|  | mCherry | Red fluorescent protein |  |
|  | tagBFP-CAAX | Blue fluorescent protein, membrane targeted |  |
|  | Venus-CAAX | Yellow fluorescent protein (YFP), membrane targeted | Fig. 2d |
|  | GCamp6f | Green fluorescent calcium indicator | Fig. S5 |
|  | Chrimson-tdTomato | Chrimson-tdTomato fusion | Fig. S5 |
|  | Jaws-mRuby | Jaws-mRuby fusion |  |
|  | TVA-mCherry | TVA-mCherry fusion | Fig. 2e, Fig 4, Fig. S3, Fig. S4, Fig. S9 |
|  | zoSADG | glycoprotein for rabies tracing | Fig4, Fig5, Fig. S9 |
|  | TeNT-GFP | tetanus toxin light chain fused with GFP | Movie 4 |
|  | DT(Diphtheria toxin) | diphtheria toxin |  |

**Table 2.**
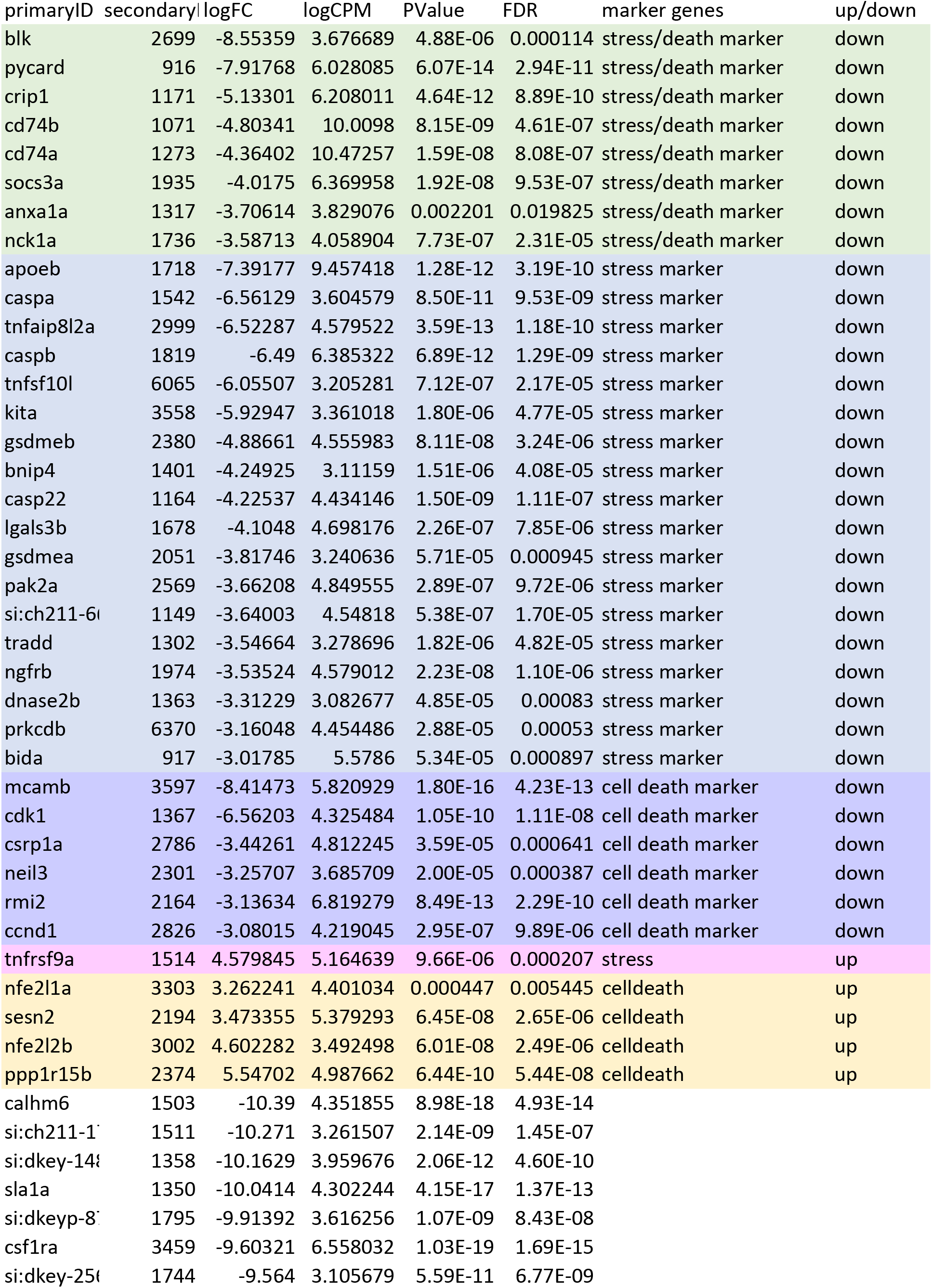
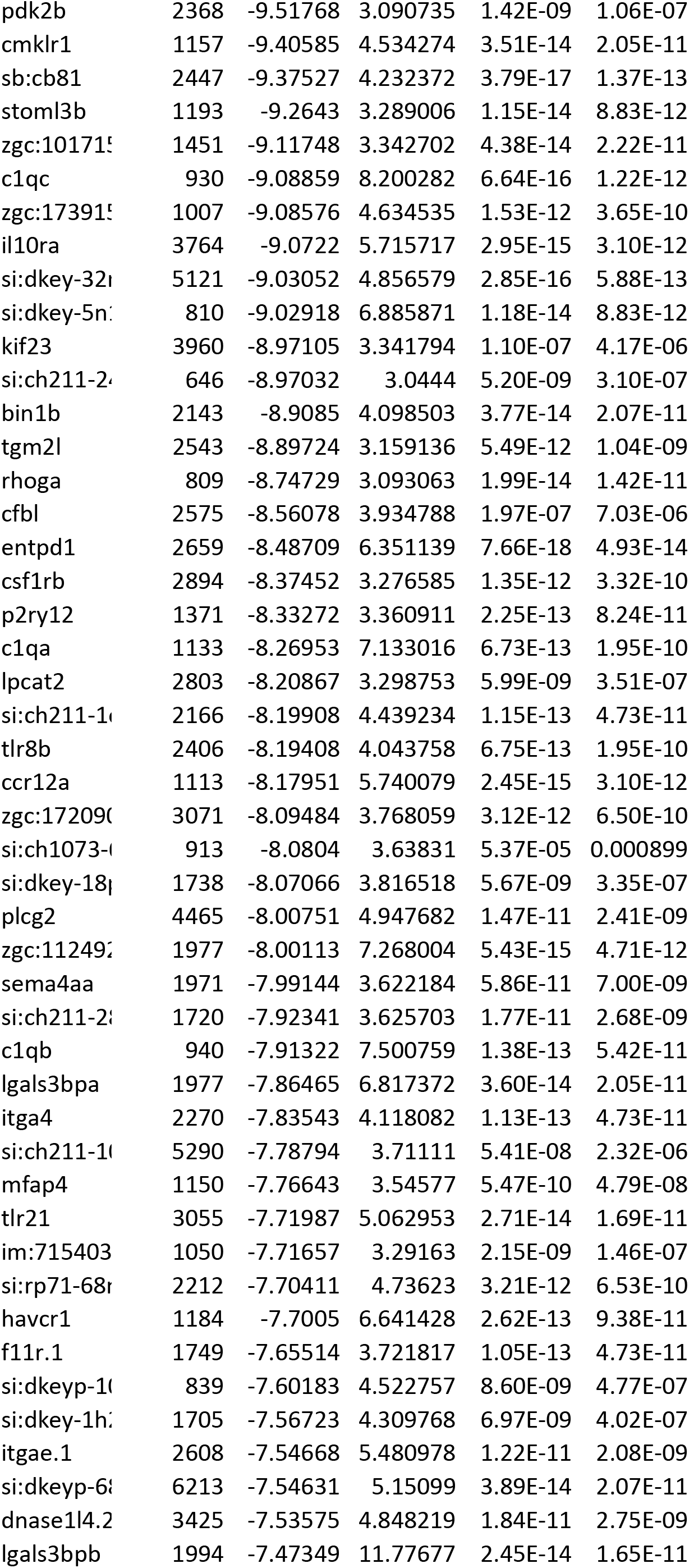
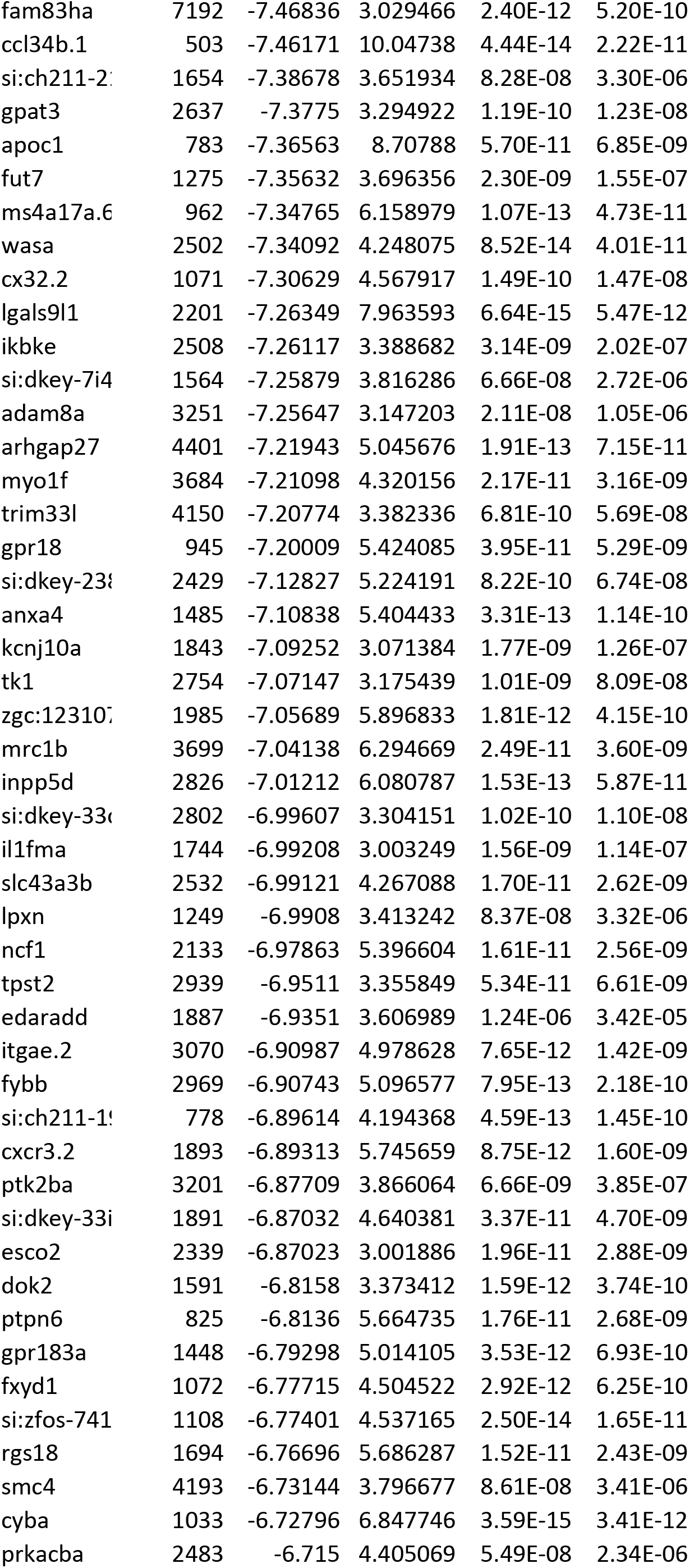
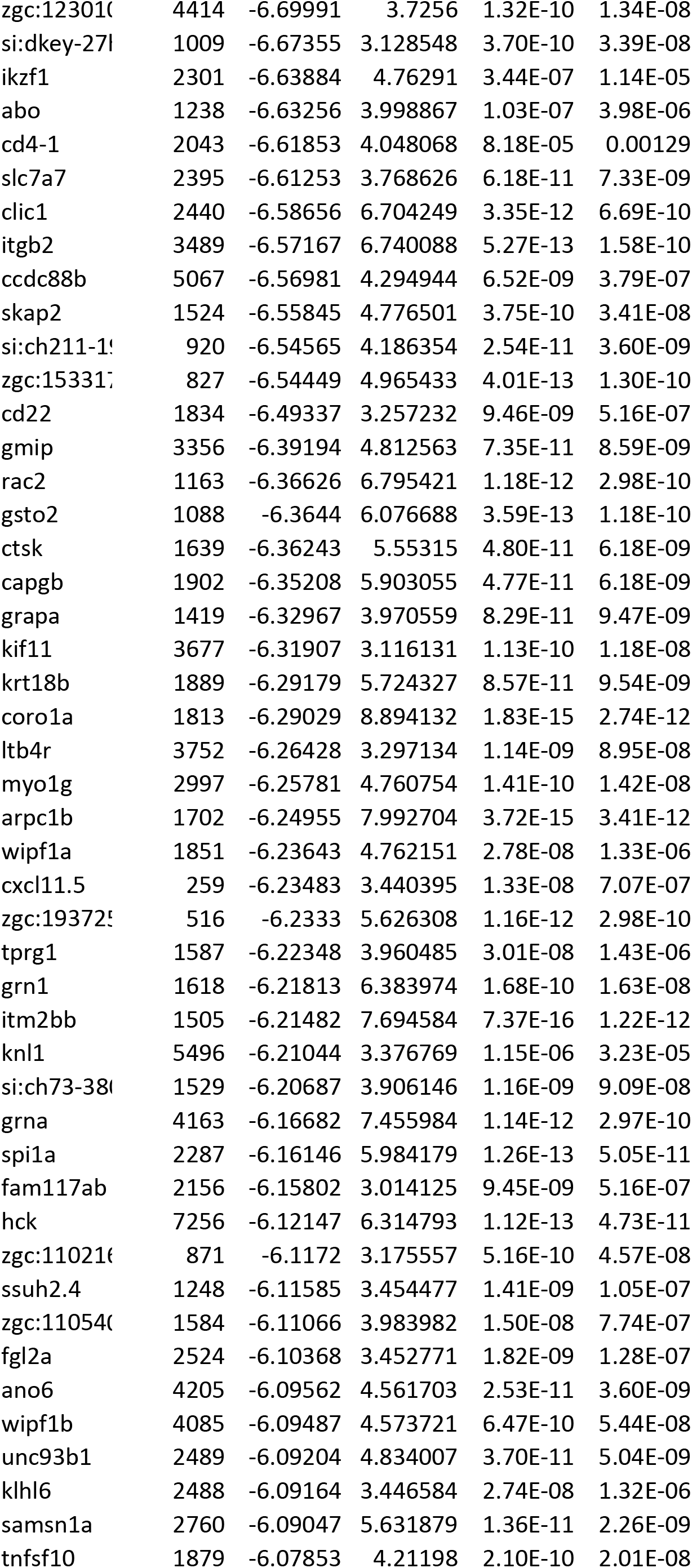
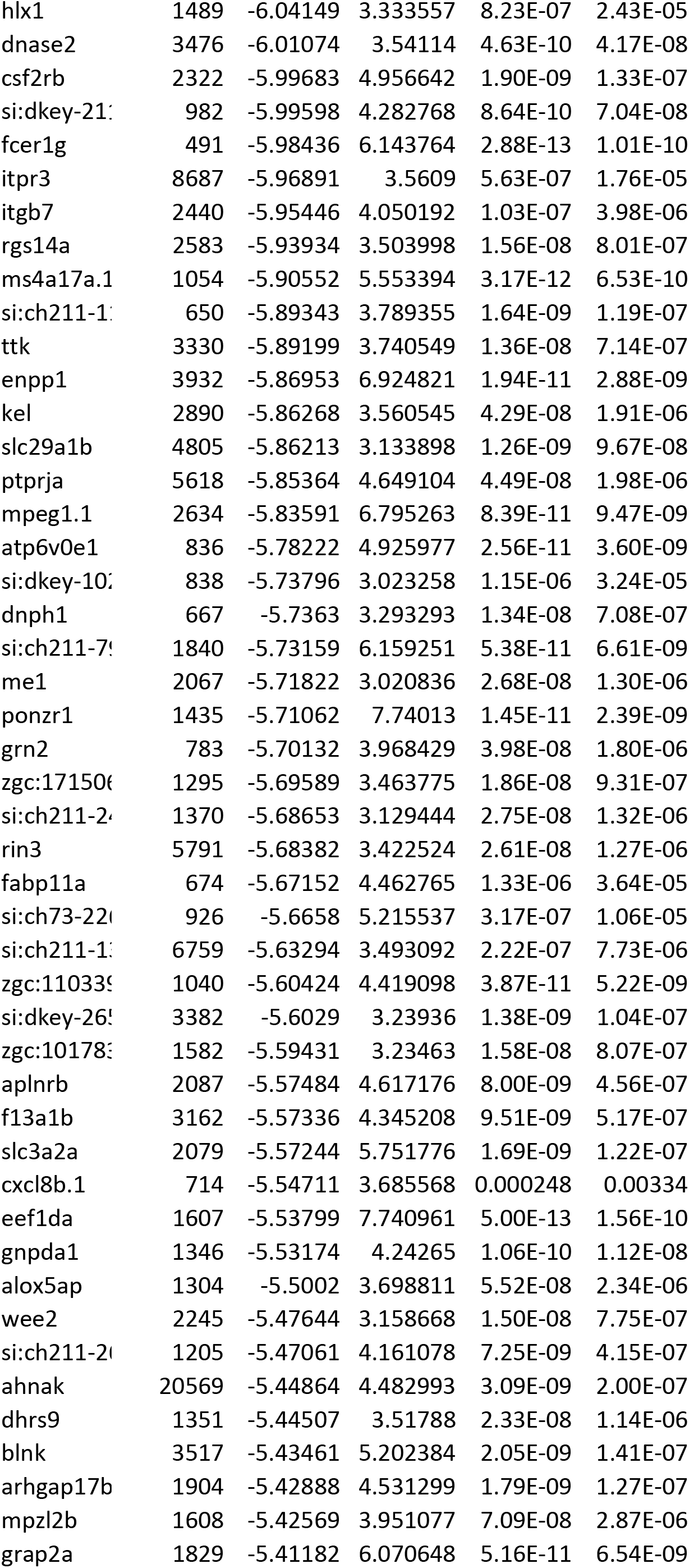
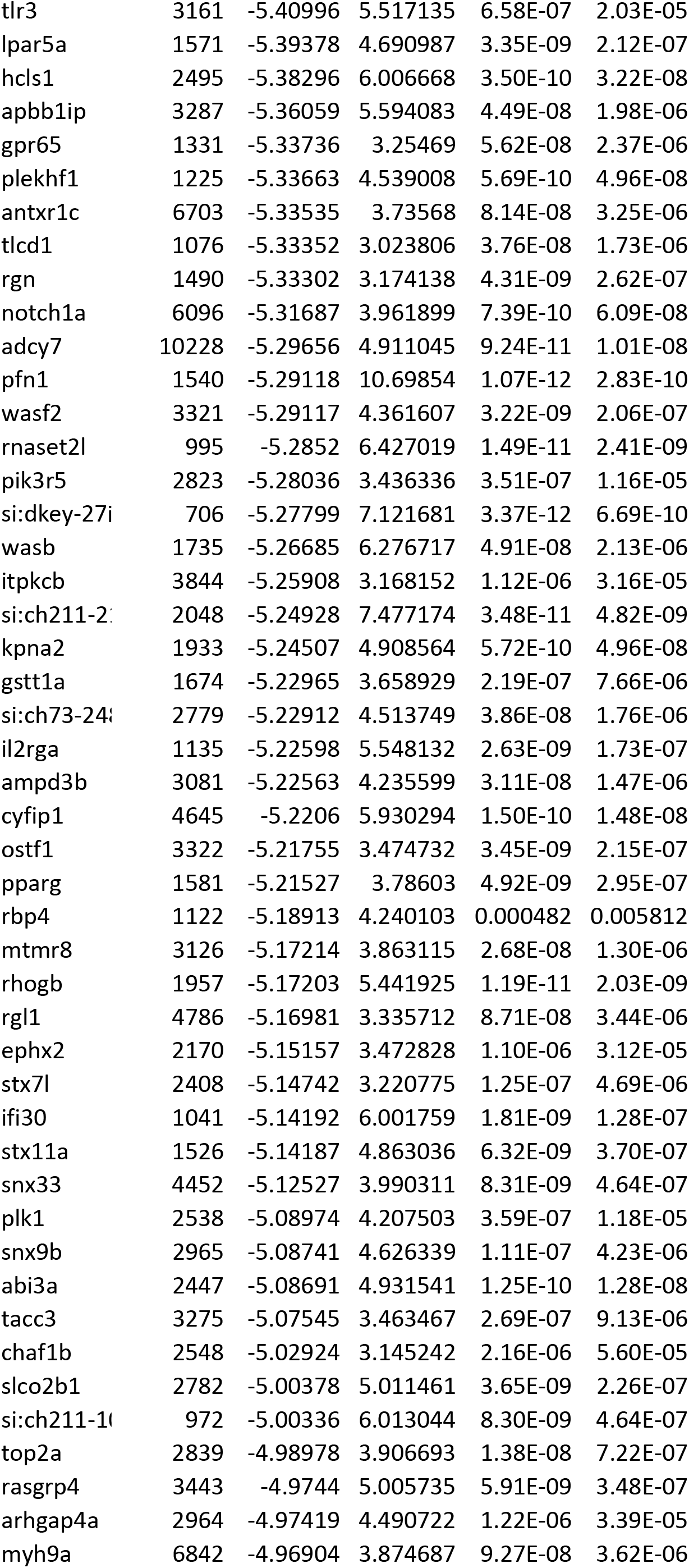
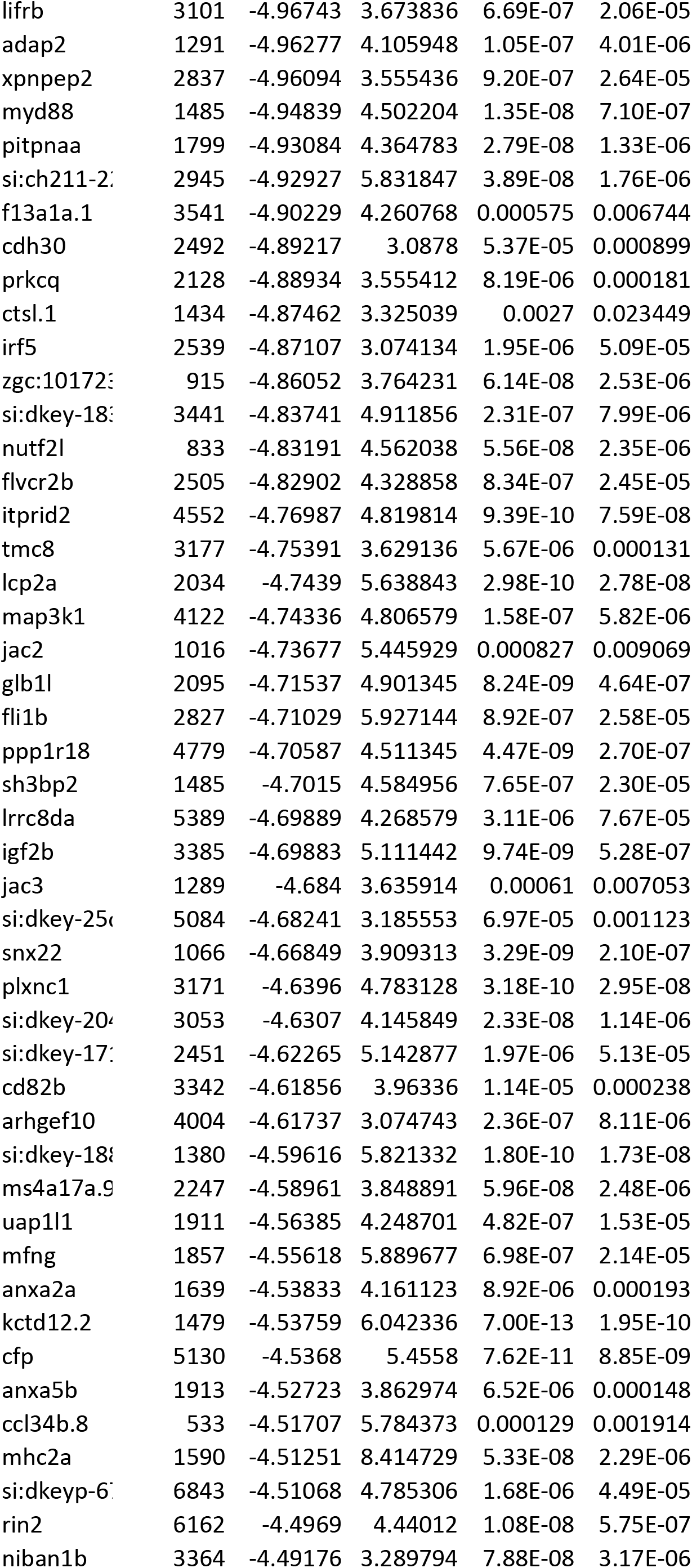
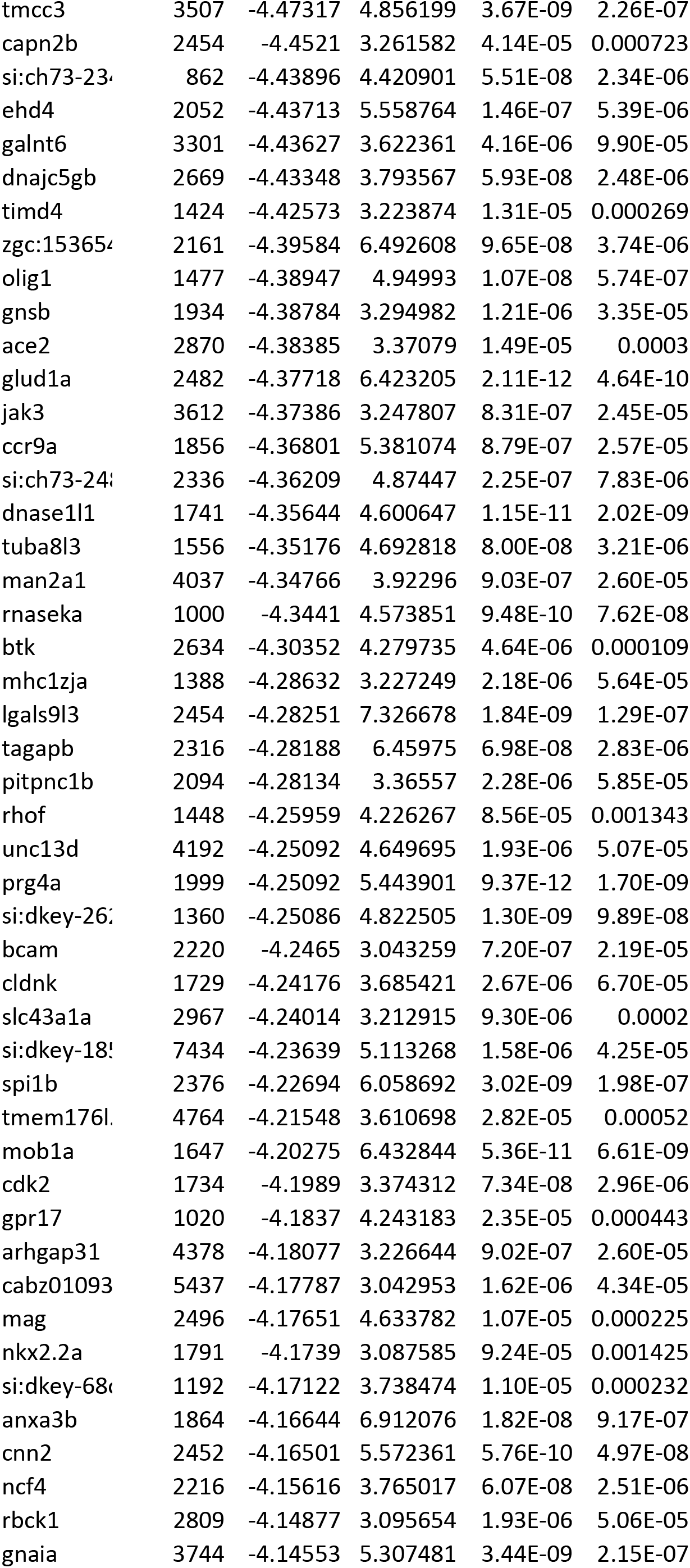
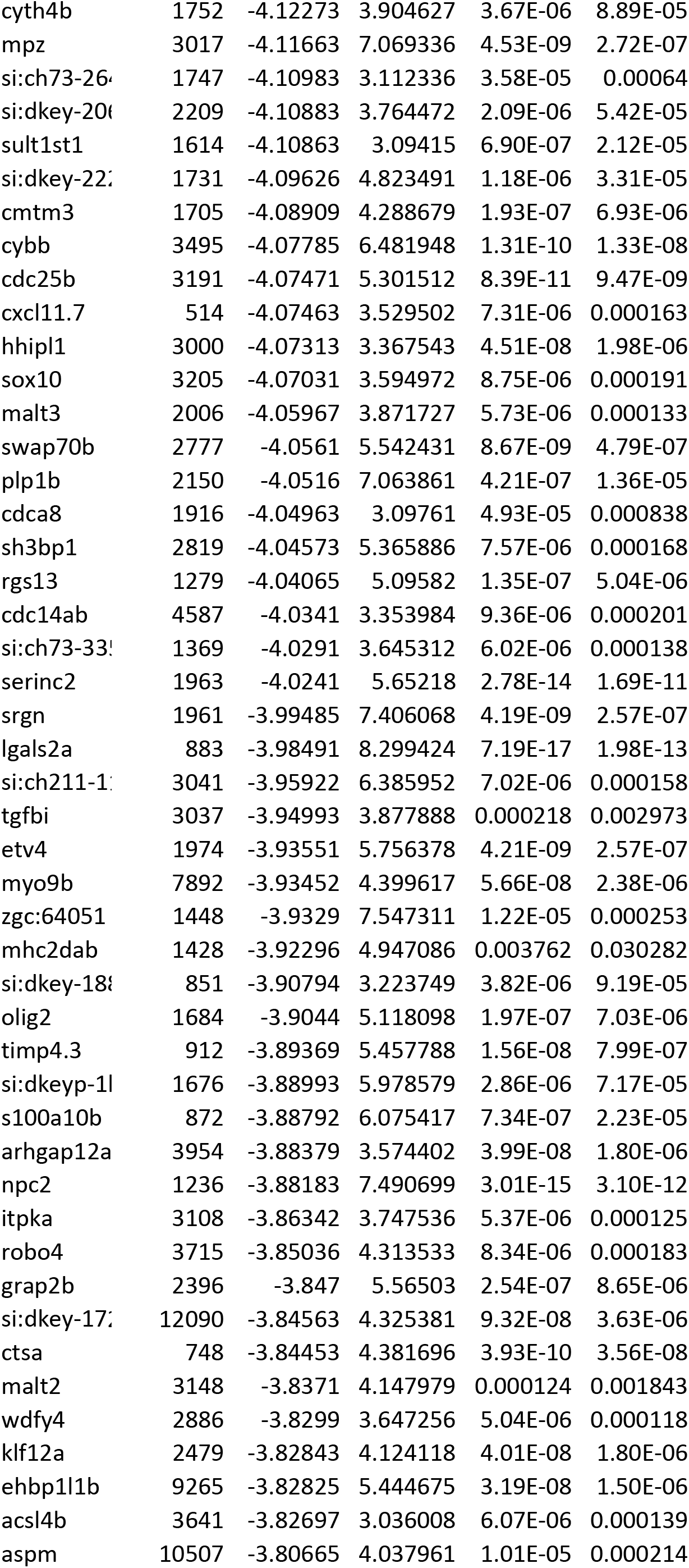
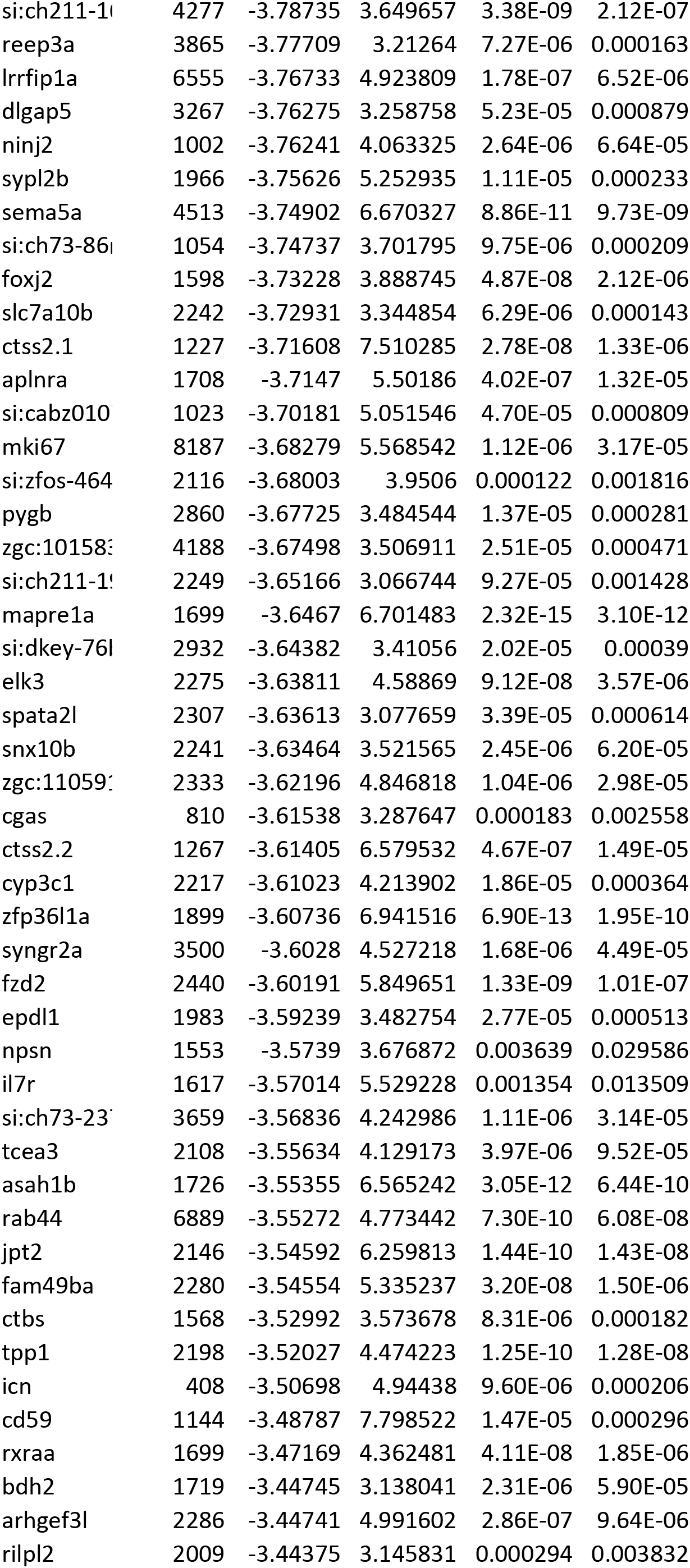
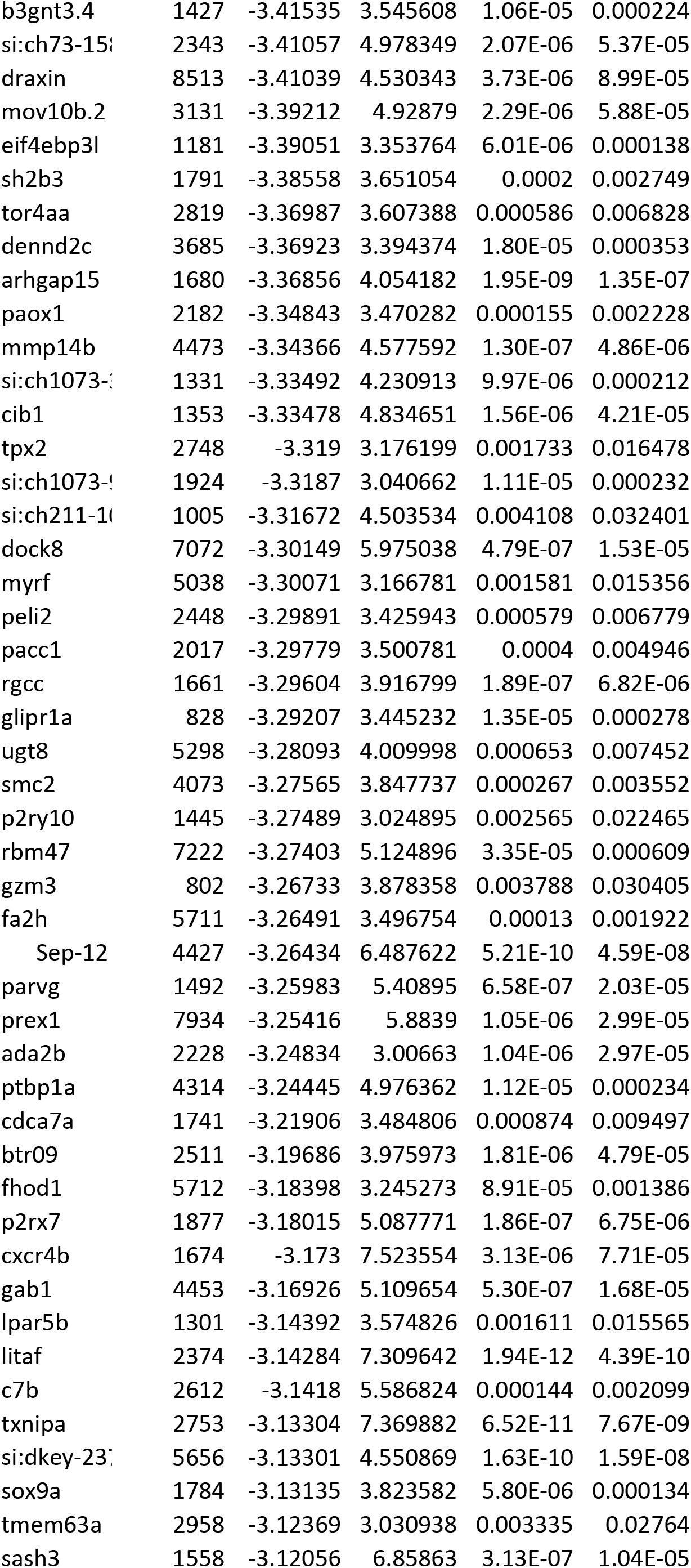
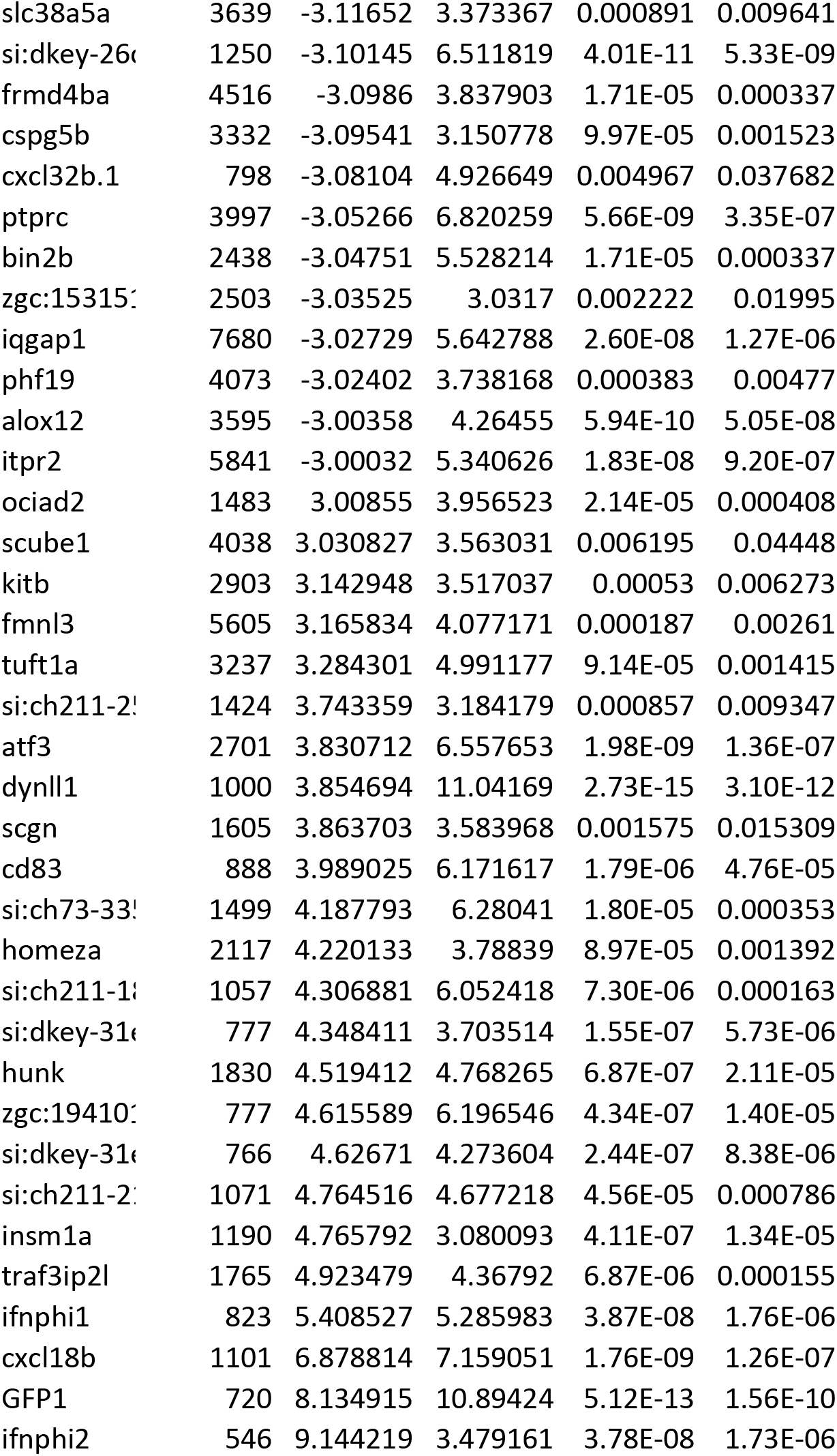
List of genes that were significantly up- or downregulated in cells infected by RVΔG. Genes associated with GO terms cellular response to stress (GO:0033554) and cell death (GO: 0008219) are listed at the top (colored rows).

**Table 3.**
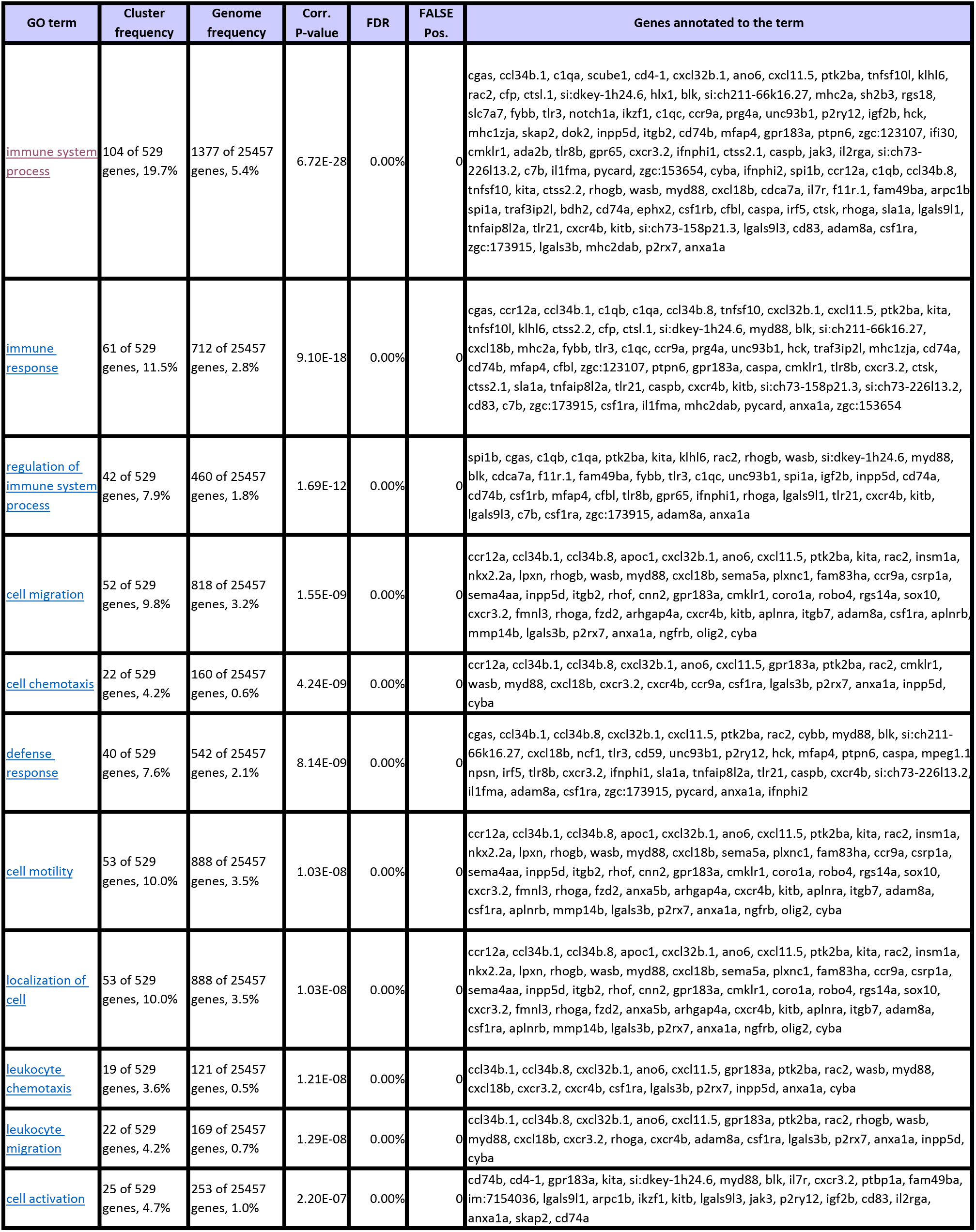
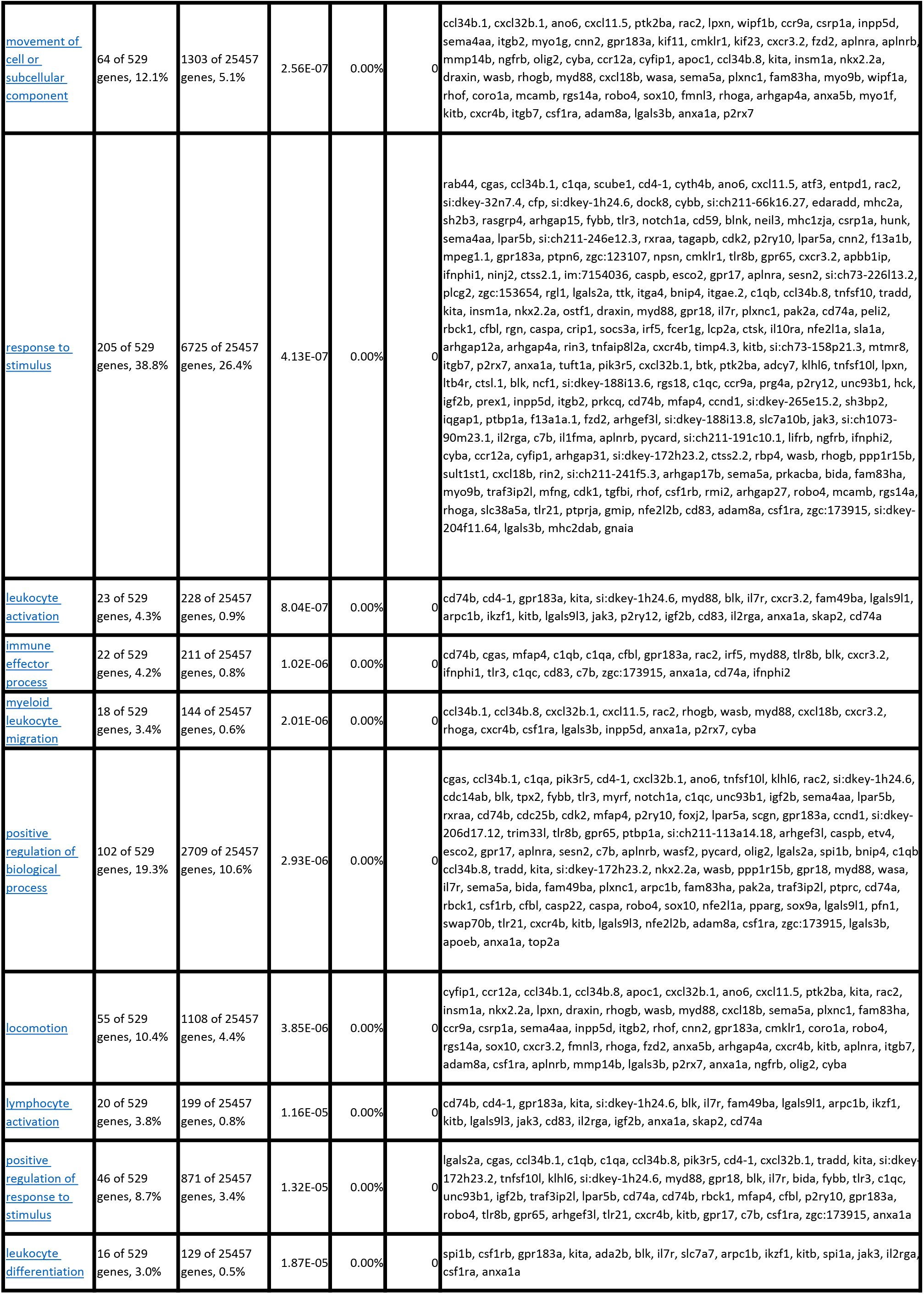
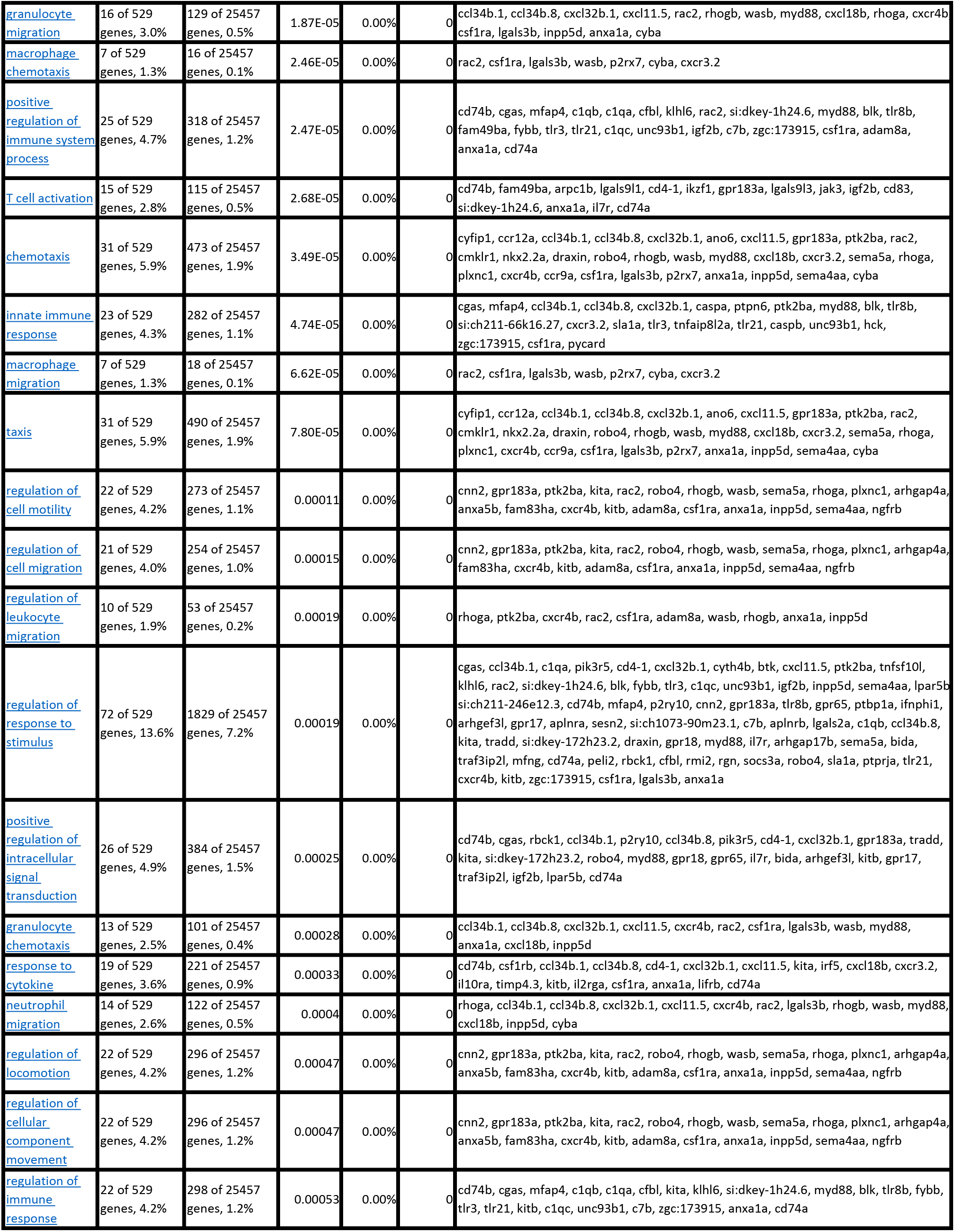
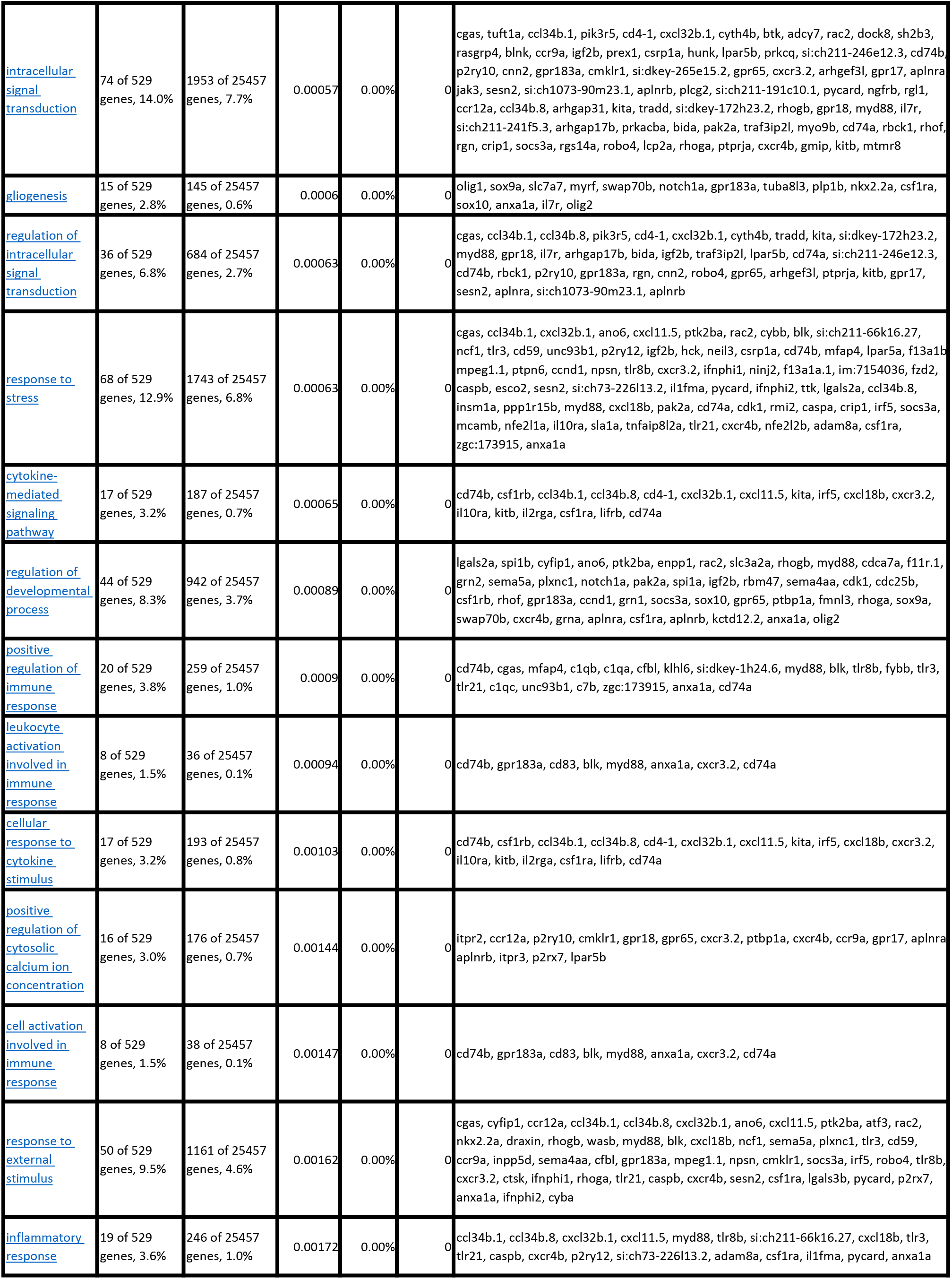
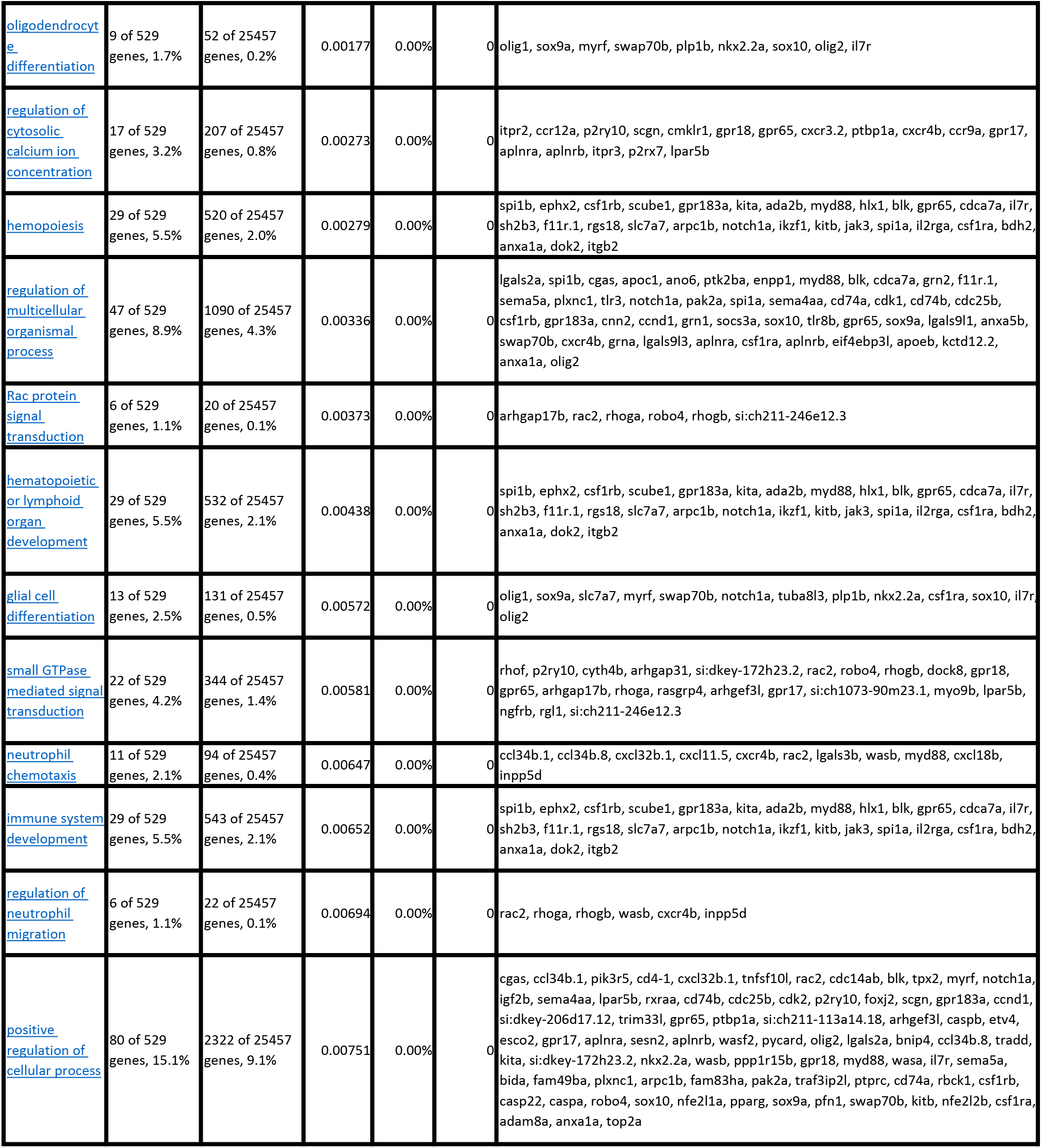
Gene ontology (GO) terms that showed a significant association with the set of regulated genes in RVΔG-infected cells (Supplementary Table 2), sorted by probability (p-value). Note that most GO terms are linked to immunity. GO terms linked to stress, cell death or synaptic functions are rare or absent.

